# Persistent cortical excitatory neuron dysregulation in adult *Chd8* haploinsufficient mice

**DOI:** 10.1101/2025.06.04.657776

**Authors:** Cesar P. Canales, Stephanie A. Lozano, Nicholas A. Frost, Karol Cichewicz, Wellington Amaral, Nicolas Seban, Ethan Fenton, Ayanna Wade, Nickolas Chu, Emily Smith, Cory Ardekani, Samuel Frank, Jeffrey Bennett, Pierre Lavenex, Aspen Kopley-Smith, Darlene Rahbarian, Melissa Corea, Daniela Perla, Liam Davis, Jiyuan Zhu, Rebecca Ortiz, Paris Beauregard, Sarah Morse, Jacob Baker, Jingqi Sun, Boxuan Ma, Ju Lu, Vikaas S. Sohal, David G. Amaral, Yi Zuo, Alex S. Nord

## Abstract

*CHD8* mutations cause autism spectrum disorder, cognitive deficits, and macrocephaly. *Chd8^+/-^* mouse models exhibit macrocephaly and transcriptional pathology, with inconsistent findings regarding neurogenesis, neuron function, and behavior. Via stereology and single nucleus transcriptomics (snRNA-seq), we found increased *Chd8^+/-^* cortical volume was not explained by increase in neuron number. Differential expression (DE) was present across cortical cell types, with excitatory neurons exhibiting high DE burden and shared and subclass-specific DE signatures. Bulk RNA-seq DE of constitutive *Chd8^+/-^* and conditional *Camk2a-Cre Chd8^+/-^*mice identified shared transcriptional pathology. DE in synaptosomal versus nuclear mRNA identified overlapping DEGs, but also significant differences and exaggerated synaptosomal changes. Building on DE findings implicating glutamatergic neurons, we found *Chd8^+/-^*mice exhibited altered excitatory neuron spine density and dynamics, decreased GCaMP activity correlation, and sleep perturbation. Thus, *Chd8* haploinsufficiency causes lasting excitatory neuron dysfunction, perturbs RNA regulation beyond transcription, and impacts neuronal properties, cortical microcircuits, and behavior.

## Introduction

Mutations in a growing list of genes cause Neurodevelopmental Disorders (NDDs) with overlapping but complex presentation, including autism spectrum disorder (ASD), intellectual disability (ID), and behavior or psychiatric disorders^1, 2^. *CHD8* encodes the chromatin remodeling factor (CRF) Chromodomain Helicase DNA-binding protein 8, and has among the highest rates of protein disrupting rare and de novo mutations in ASD cases^1^. CHD8-associated NDD (*CHD8*-NDD) has core but variably expressed features including ASD, ID, and macrocephaly, as well as sleep, gastrointestinal, psychiatric, and neurological issues^3–9^. Clarifying the links between *CHD8* haploinsufficiency, macrocephaly, and pathology may offer critical insights into the etiology of the 15% of ASD patients that exhibit more severe cognitive and behavioral phenotypes often with poorer prognosis^10^. Furthermore, CHD8 has been identified as a central regulator in network studies of ASD-associated loci^11–13^, and serves as a key example of the many CRF genes implicated in ASD and other NDDs^14–18^. Defining the pathological consequences associated with *CHD8* haploinsufficiency is thus an important step towards understanding both *CHD8*-specific mechanisms and broader NDD neurobiology.

*Chd8* is ubiquitously expressed and knockout is embryonic lethal in mice^19, 20^. NDD-relevant phenotypes have been observed across studies of heterozygous *Chd8^+/-^*mouse models^20–26^. Among the consistent findings across constitutive *Chd8^+/-^* mouse models is the presence of adult macrocephaly and transcriptional pathology^20, 22, 24, 25^. Bulk RNA-seq on developing and adult *Chd8^+/-^* mouse brain has been performed in a number of studies, finding relatively subtle effects but replicated perturbation of energetics and protein homeostasis and downregulation of synaptic and neuronal genes^20, 21, 24, 25, 27–30^. Recent single cell studies of *Chd8^+/-^* mice suggest subtle changes in neurodevelopment and perturbations to neurons^31–33^, but have lacked resolution to analyze specific subtypes in mature cortex. Studies of *Chd8^+/-^* mouse behavior and cognition have been more variable^20–26^, with evidence that some variability is due to genetic background^34, 35^. Multiple studies of *Chd8* haploinsufficiency have identified changes to electrophysiology in the cortex and in in vitro^21, 24, 36, 37^, indicating impacts on neuron function. In vitro studies in human and non-human primate neurons and human cortical organoids also indicate perturbation to neurogenesis, increases in organoid size paralleling macrocephaly, and neuronal electrophysiological and network level dysfunction^31, 32, 38–41^. Finally, conditional *Chd8* knockout studies have been conducted targeting specific cell types such as excitatory neurons^36, 42–46^, oligodendrocytes^47–49^, and astrocytes and microglia^50, 51^. Many of these studies have primarily focused on homozygous conditional mutants, which may not accurately model the heterozygous mutations relevant to NDDs.

Despite extensive study of *Chd8* as a representative ASD/NDD risk gene, key questions remain regarding how heterozygous mutation leads to pathology at the cellular, molecular, and circuit level. First, no published studies have linked adult macrocephaly to changes in cellular composition or described cell-type specific molecular pathology in adult *Chd8^+/-^*mouse brain. Second, it remains unclear whether neuronal dysfunction has arisen from developmental disruption or reflects a continued requirement for full *Chd8* dosage in mature neurons, and whether reported RNA expression changes are due to transcriptional or post-transcriptional mechanisms. Third, circuit-level consequences to behavior beyond sociability and cognition remain largely unexplored. To address these gaps, we characterized adult *Chd8^+/-^* mice and cerebral cortex across cellular, molecular, functional, and behavioral domains (**Figure 1A**). Our findings reveal selective vulnerability of cortical glutamatergic neurons to *Chd8* haploinsufficiency, implicate disrupted post-transcriptional regulation of synaptosomal RNA homeostasis, and demonstrate associated changes in dendritic spine dynamics, impaired neuronal signaling coherence, and altered sleep architecture in *Chd8^+/-^*mice.

**Figure 1.**
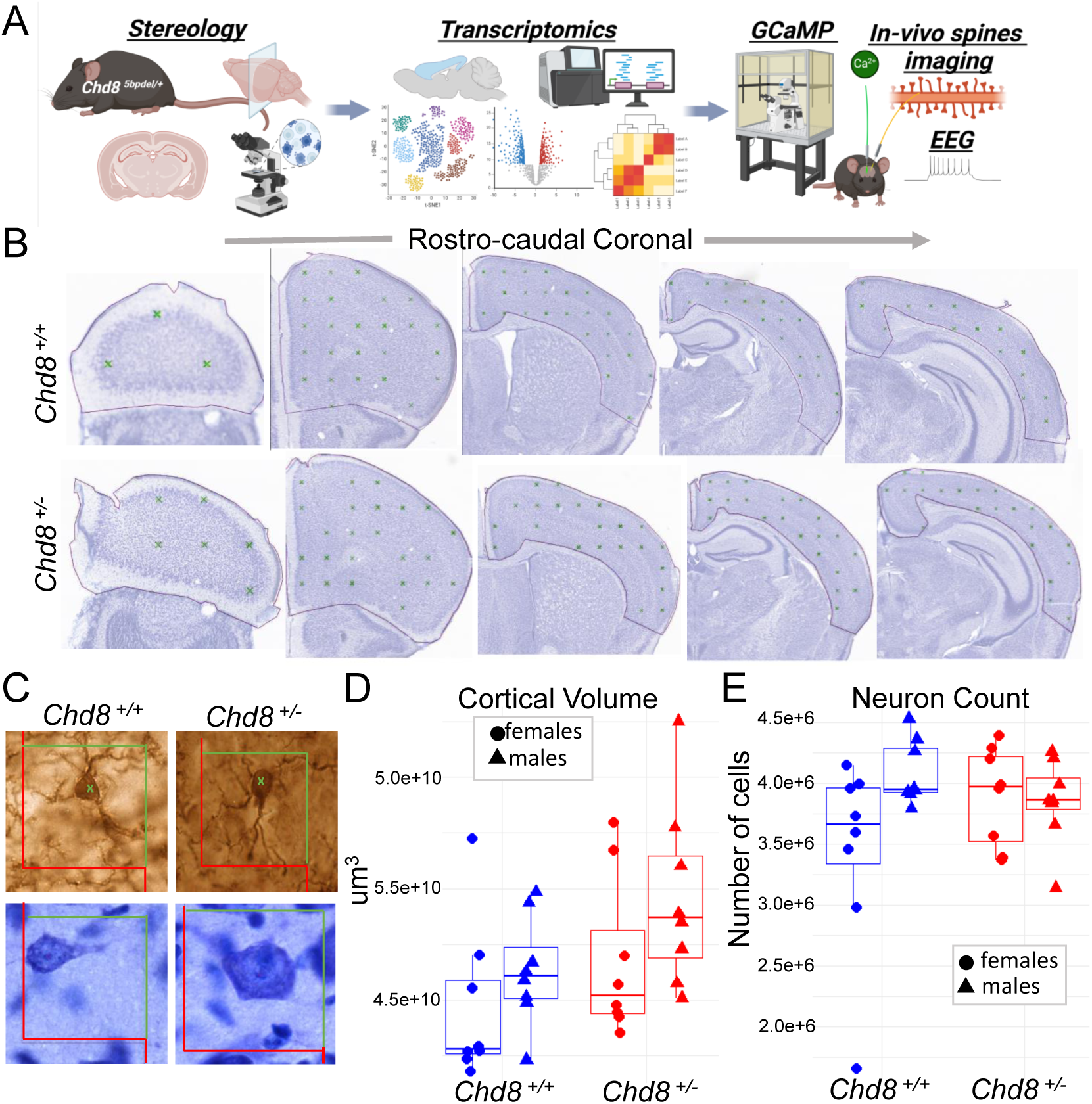
Increased cortical volume in *Chd8^+/-^* mice without a corresponding increase in neuron number. (A) Schematic overview of study investigating the role of *Chd8* haploinsufficiency in mature mouse cortex. (B–C) Representative coronal sections across the rostro-caudal extent of the cerebral cortex used for unbiased stereological analysis. (B) Cortical boundaries as determined for volume measurements. (C) Cellular organization visualized using Nissl staining (top) and NeuN immunostaining (bottom) in *Chd8^+/+^* and *Chd8^+/−^* mice. (D–E) Stereological profiles across the Z-plane showing (D) cortical volume estimates based on the Cavalieri method and (E) total neuron number as assessed by unbiased stereology in the same rostro-caudal regions. Data are stratified by sex and genotype. Error bars represent SEM. Statistical comparisons were performed using two-way ANOVA (Cortical volume: P=0.032 unstratified, P=0.009 males; P=0.61 females; Neuron Count: P=0.76, unstratified; P=0.37, males; P=0.30, females Student’s t-test, N = 8 per sex/genotype).

## Results

### Description of Chd8 mutant mouse lines

Constitutive mutant *Chd8* mice were generated on the C57BL/6N background via CRISPR/Cas9 targeting exon 5, resulting in a 5 bp deletion that leads to reduced *Chd8* mRNA and protein and is early embryonic homozygous lethal. We previously characterized heterozygous mutants from this line (referred to as *Chd8^+/5bpdel^,* herein *Chd8^+/-^*), reporting differential expression and splicing, altered cortical proliferation, increased brain volume, and phenotypes of altered exploration in open field assay and deficits in fear conditioning and novel object recognition^20^. We also use a conditional *Chd8* loxP mutant on the C57BL/6J background generated by Jackson Laboratory where exon 3 is ablated in the presence of CRE (*Chd8^em3Lutzy^*). These were mated to *Camk2a*-CRE (B6.Cg-Tg(Camk2a-cre)T29-1Stl/J; JAX strain: 005359) to direct conditional heterozygous ablation in post-mitotic forebrain glutamatergic neurons in this line, herein referred to as *Chd8^+/flox^*.

### Increased adult cortical volume without increase in neuron number or soma size in Chd8^+/-^ mice

We previously identified increased cortical volume and macrocephaly in *Chd8^+/-^* mice via structural MRI^20^. Here, we aimed to replicate this finding and determine whether the volume increase results from a higher number of neurons. We performed unbiased stereological analysis on Nissl-stained brain sections from male and female heterozygous *Chd8^+/-^* mice and wild-type littermates at postnatal day 42 (P42) (**Figure 1B-C**), an age marking the end of adolescence and a mature stage of mouse brain development^52^. We examined 15–18 representative coronal sections per mouse using established stereological methods to accurately assess cortical volume, neuronal distribution, and soma size. Cortical volume was estimated using the Cavalieri method^53, 54^, revealing a significant overall increase in *Chd8^+/-^* mice compared to wild-type littermates (**Figure 1D**, P=0.032, Student’s t-test). When stratified by sex, the increase in cortical volume was more pronounced and independently significant in males (P=0.009 males; P=0.61 females, Student’s t-test). Comparing volume across rostral-caudal sections, *Chd8^+/-^* males showed increase in more rostral regions (**Figure S1A**).

Next, we investigated whether the cortical volume expansion was associated with an increase in neuronal numbers or soma size. Using the optical fractionator method^55^, we estimated neuron counts across the cortex. We did not find significant differences in total neuron number in either the full cohort or sex-stratified analyses (P=0.76, unstratified; P=0.37, males; P=0.3, females) (**Figure 1E**). Finally, we assessed neuronal soma size and found no significant differences between genotype groups in full or sex-stratified analyses (P=0.23, unstratified; P=0.81, males; P=0.19, females) (**Figure S1B**). Taken together, these findings replicate the cortical volume increase observed in *Chd8^+/-^* mice, with a stronger effect in males, and demonstrate that this expansion is not explained by an increase in neuronal number or soma size. Instead, the increased cortical volume without an increase in numbers results in decreased glutamatergic neuron density in the *Chd8^+/-^* cortex.

### Cell-type specific cellular and molecular pathology in Chd8^+/-^ P60 cortex

Towards characterizing differences in RNA expression associated with *Chd8* haploinsufficiency in adult (P60) mouse cortex, we performed single nucleus RNA sequencing (snRNA-seq) on constitutive *Chd8^+/-^* mice, as well as bulk RNA-seq on *Chd8^+/-^* and conditional *Camk2a-CRE Chd8^+/flox^* mutants, and for nuclear and synaptosomal RNA from *Chd8^+/-^*mutants (**Figure 2A**). For snRNA-seq, we analyzed *Chd8^+/-^* (male n = 3, female n= 2) and WT littermates (male n=2, female n=3 WT). Following quality control and ambient RNA correction^56^ (**Figure S2A**), there were 11,751 and 12,169 nuclei from *Chd8^+/-^* and WT samples, respectively, which we mapped to 18 cortical cell types based on marker gene expression and integration with the Brain Initiative Cell Census Network mouse brain atlas^57^ (see methods) (**Figure 2B** and **Figure S2B**). There was balanced representation of male and female samples and *Chd8^+/-^* and WT samples (**Figure 2C**). Expected major cortical cell types were identified as shown by expression of marker genes (**Figure 2D**). Cell identity was split at three levels: cell type 3 representing general classes (e.g. glutamatergic neurons), cell type 2 associated with specific canonical subtypes (e.g. L4/5 IT), and cell type 1 representing data-driven further sub-clustering of subtypes (e.g. L4/5 IT Glut1). Following UMAP generation and cell type identification, we tested for differences in the relative representation of different cell types, comparing cell type proportions between *Chd8^+/-^* and WT groups (**Figure 2E**). At the cell type 2 level, there was significant increase in OPC and decrease in L6 IT, with trending increase in astrocytes and decrease in microglia. These findings are consistent with the results from stereology, demonstrating that macrocephaly and increased cortical volume in *Chd8^+/-^* mice is not due to increase in number of glutamatergic neurons and further suggesting that increased cortical volume is not explained by expansion of specific cell types.

**Figure 2:**
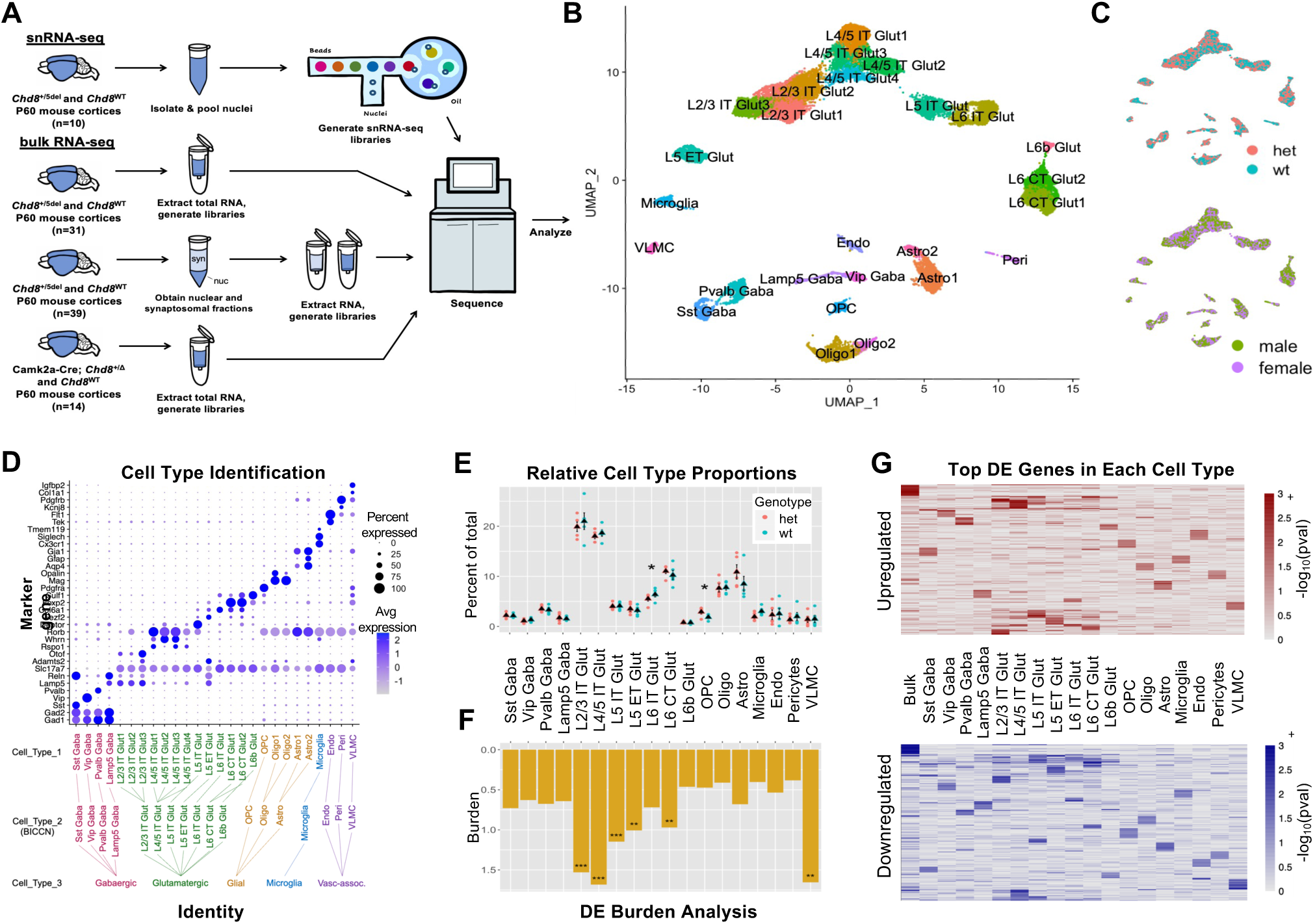
Cell type-specific pathology in adult *Chd8^+/-^* mouse cortex. A) Schematic depicting experimental design for RNA-seq and snRNA-seq experiments. B) UMAP of 23,920 cortical nuclei from P60 *Chd8^+/-^* and WT littermates with cell types annotated. C) Clustering of *Chd8*^+/-^ (HET) and WT nuclei (top) and male and female nuclei (bottom) in UMAP space. D) Cell type identification dot plot representing average expression of cell type-specific marker genes in each cell type. Tree (bottom) shows 3 levels of cell type identification (Cell_Type_1-3). E) Cell type proportion comparison between *Chd8^+/-^* (HET) and WT nuclei, revealing a significant decrease in mutant L6 IT Glut nuclei (*p = 0.027) and increase in mutant OPCs (*p = 0.048) (Student’s t-test). F) DEG burden across cell types (Burden = DE genes / expressed genes * 100). Significant DE burden was found in L2/3 IT (*p = 0.00001), L4/5 IT (*p = 0.00001), L5 IT (*p = 0.00001), L5 ET (*p = 0.0045), L6 CT Glut (*p = 0.0004), and VLMC (*p = 0.0009) nuclei (permutation test, n= 10,000). G) Heatmap showing –log10(pval) of top 25 unique upregulated (top) and downregulated (bottom) bulk and pseudobulk DEGs in each cell type (p < 0.1).

Next, we performed cell-type specific differential expression (DE) analysis using a cluster-level pseudobulk framework^58^, where reads are from the relevant cell type are aggregated at the sample level, followed by DE testing using DESeq2^59^ with experimental batch included as a covariate. We performed DE analysis at each level of cell type identity (**Tables S5-S7**), with a focus on cell type 2 for results here. Comparing DE burden (defined as the number of total DE genes divided by genes passing expression criteria for testing), glutamatergic cell types along with VLMC showed the largest impacts (**Figure 3F**). Among glutamatergic neurons, L2/3 intra-telencephalic (IT) and L4/5 IT neurons had the highest burden. DE results and burden were stable using either the full dataset or when down-sampling for equivalent cell/nuclei numbers across cell types and when using both uncorrected and ambient RNA corrected expression values, and similar differential expression effect direction and sizes were present in stratified male and female samples (**Figure S2C-E**). At the single gene level, the top DE genes (DEGs) varied by cell type, with the exception that glutamatergic neuron subtypes, showed considerable overlap in effects (**Figure 3G**).

**Figure 3:**
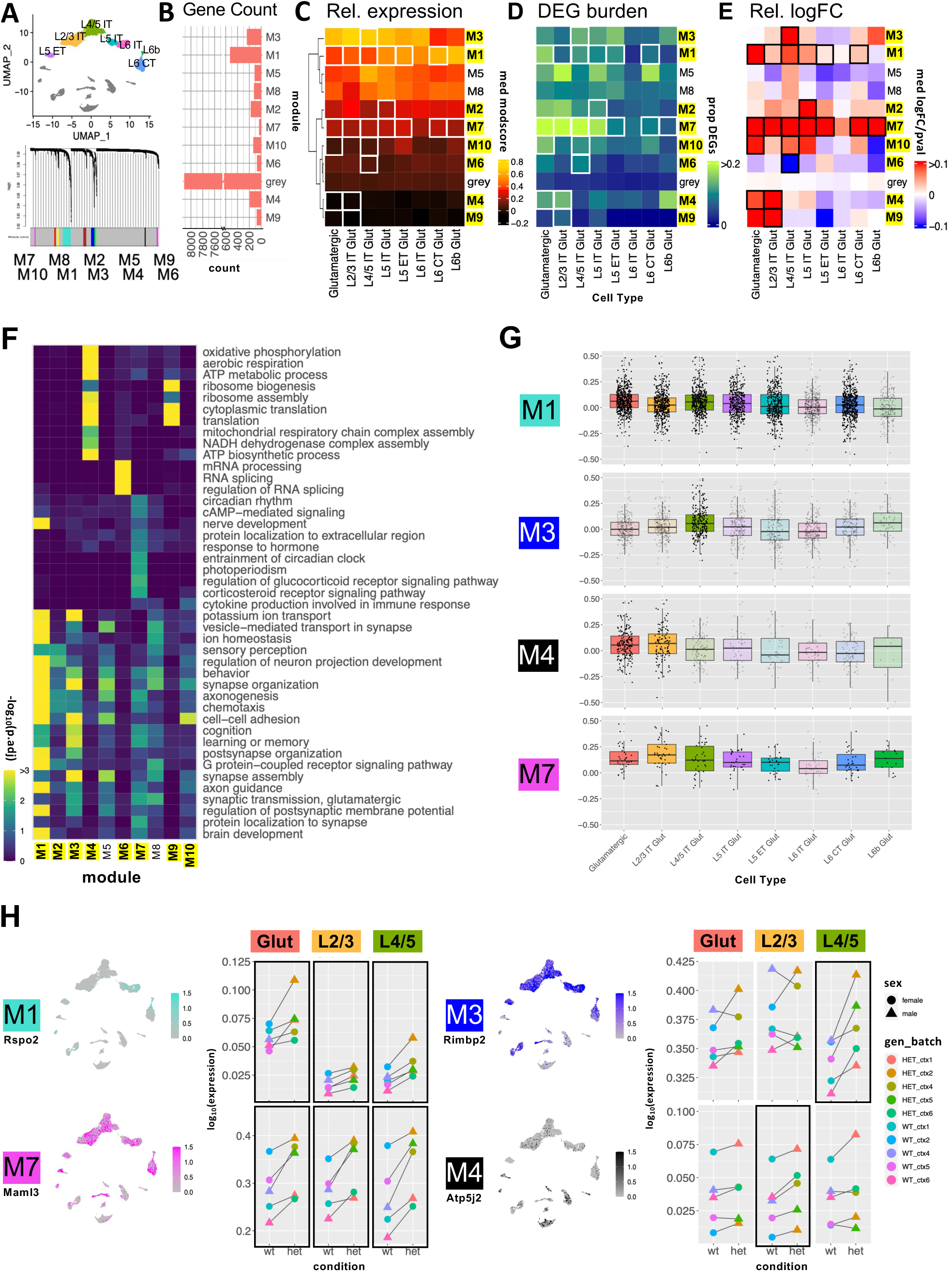
hdWGCNA reveals differentially expressed glutamatergic cell coexpression modules. **A)** 10 glutamatergic modules identified via hdWGCNA B) Number of genes per module. C) Relative expression of glutamatergic modules across glutamatergic neuron subtypes (module score = average expression of all module genes compared to expression-matched background genes; DEMs identified via comparison of median pseudo-bulk logFC of module genes to that of the grey module via t-test; DEMs (*p < 0.05) in at least one population are labeled in yellow. D) DEG burden (calculated as in Fig. 2F) of each DEM in each cell type E) Heatmap of relative median pseudo-bulk logFC of module genes for glutamatergic cell types scaled by p-value. White and black boxes in C-E indicate the cell types in which each corresponding module was DE. F) Representative significant GO terms for Biological Processes enriched in glutamatergic modules. (*p < 0.05, **p < 0.01, ***p < 0.001). G) Box plots for 4 representative DEMs showing logFCs of module genes in each glutamatergic subtype. Non-opaque boxes indicate cell types where the module shown was significantly differentially expressed (P-value < 0.05). H) Feature plots (left) showing relative expression of representative genes from DEMs of interest in UMAP space. Dot plots (right) show median sample-level expression of the same genes. Black boxes indicate cell types where the corresponding module is differentially expressed, or where the direction of change in module expression is consistent across batches.

We performed gene set enrichment analysis (GSEA) based on DE rank using the clusterProfiler package^60^ (**Figure S2F**). Only glutamatergic neuron subtypes had DE signatures strongly associated with GO terms (FDR < 0.05), with the strongest signatures in L2/3 IT and L4/5 IT. Using a relaxed threshold of uncorrected P < 0.05, upregulated genes in glutamatergic neurons showed shared enrichment for pathways including *Synapse organization*, *Synaptic signaling*, and *Axon guidance*, and *Neuron differentiation*. Finally, we applied NeuronChat^61^ to explore cell-cell communication networks comparing receptor-ligand relationships in *Chd8^+/-^* and WT cortex based on snRNA-seq data (**Figure S3**). This identified reduced Neurexin-Neuroligin interaction strength in *Cdh8^+/-^* mice, with the strongest effect in L2/3 IT neurons (**Figure S3H-K**). At the level of information flow for individual receptor-ligand pairs, NeuronChat identified a trend of increased information flow involving glutamate receptor-ligand interactions in *Chd8^+/-^* mutants as well as decreased flow for specific Neurexin-Neuroligin pairs (**Figure S3L**). These findings highlight disrupted synaptic signaling in *Chd8^⁺/⁻^* mice and suggest that glutamatergic neuron subtypes, particularly L2/3 IT and L4/5 IT neurons, are disproportionately affected in the adult cortex.

### Shared and cell-type specific DE co-expression modules in Chd8^+/-^ glutamatergic neurons

To capture systems-level expression patterns in glutamatergic neurons, we used hdWGCNA^61^ to identify gene co-expression modules and then identified differentially expressed modules (DEMs) based on log2-transformed Fold Change (logFC) of module genes (see methods for details). All glutamatergic cell subtypes were used as input to hdWGCNA, which yielded 10 modules accounting for 1,691 (17%) genes, with the remaining 8,115 genes assigned to the grey group (**Figure 3A-B, Table S8**). 8/10 modules were DEMs in at least one glutamatergic subtype, as indicated by highlighted label and box outlines in **Figure 3C-E**. Two modules (M1 and M7) were DEMs across most glutamatergic subtypes, whereas the other six DEMs (M2, M3, M4, M6, M9, and M10) showed genotype differences in only one or two subtypes. Module level differential expression direction and effect size varied, though all but one DEM (M6 in L4/5 IT) were upregulated in *Chd8^+/-^* mutants. The modules varied in gene number (**Figure 3B**) and relative expression patterns (**Figure 3C**) across glutamatergic subtypes and cortical layers. The proportion of module genes that were individually DE (**Figure 3D, Figure S4A**) and the module-level logFC (**Figure 3E**) similarly varied across modules and subtypes.

The glutamatergic co-expression modules showed different association to biological processes based on GO enrichment (**Figure 3F**). First, 6/10 modules showed overlapping enrichment for a variety of general neuron-associated terms (e.g. *Synapse assembly*, *Axon Guidance*, *Behavior*) without strong module-specific GO terms. These included DEMs M1, M2, and M3, as well as non-DEMs M8 and M5. These modules also exhibited differential expression across glutamatergic neuron subtypes and cortical layers, suggesting that their co-expression is primarily driven by expression co-variation linked to cell identity, rather than by shared biological pathways or genotype effects. The remaining four modules (M4, M6, M7, M9) were all DEMs and featured strong module-specific GO enrichment: M4 and M9, associated with *Aerobic Respiration* and *Translation* and upregulated in L2/3 IT, M6, associated with *mRNA processing* and downregulated in L4/5 IT, and M7, which was broadly upregulated and associated with general neuronal terms as well as GO terms represented by *Circadian rhythm*. In addition to module-specific GO terms, these four modules did not show subtype- or laminar-associated differences in relative expression and had DE signatures limited to individual glutamatergic subtypes. We highlight gene-level differential expression patterns for DEMs of high interest in **Figure 3G**, focusing on M1 and M7 modules that are broadly DE across glutamatergic subtypes, and M3 and M4 modules that are subtype-specific DEMs with prominent effects in L4/5 IT and L2/3 IT, respectively. **Figure 3H** shows representative gene expression patterns from these DEMs visualized in UMAPs and pseudobulk-level relative expression plots. See **Figure S4B-C** for extended modules visualizations. M1 (featuring *Rspo2*) includes a large set of genes that are expressed most highly in deep layer excitatory neurons (L6 CT and L6b), associated with general neuronal and synaptic functions. Although it includes both up- and downregulated genes, M1 shows a significant bias towards upregulation with the strongest effects in L4/5 IT and L5 IT subtypes. In contrast, M7 (featuring *Malm3*) is a smaller module exhibiting consistent upregulation across all glutamatergic subtypes. It is associated with general neuronal function as well as circadian rhythm and hormone response pathways. M3 (featuring *Rimbp2*) is selectively upregulated in more superficial glutamatergic subtypes L4/5 IT neurons, with enrichment for neuronal and synaptic processes. In contrast M4 (featuring *Atp5j2*) shows low expression across glutamatergic subtypes and GABAergic neurons, but relatively higher expression in non-neuronal cell types, with gene ontology terms related to translation and mitochondrial ATP metabolism. The shared dysregulation of M1 and M7 DEMs mirrors overlapping GO enrichment patterns across glutamatergic subtypes observed in GSEA (**Figure S2F**), supporting convergent molecular signature across glutamatergic neurons. The subtype-specific dysregulation of M3 and M4 suggest genotype-dependent perturbations of distinct biological processes particularly affecting L2/3 IT and L4/5 IT neuron populations.

### Bulk RNA-seq comparison identifies shared signatures in constitutive versus conditional Chd8 mutants but divergent effects in nuclear versus synaptosomal mRNA fractions

We employed bulk RNA-seq as a complementary approach to snRNA-seq, assaying DE signatures across four contrasts: mRNA from P60 cortex from constitutive *Chd8^+/-^* and from conditional *Camk2a- Cre* x *Chd8^+/flox^* mice, and compartment-enriched nuclear and synaptosomal mRNA from P60 cortex from *Chd8^+/-^* mice (**Figure 4**). See **Tables S9-S11** for sample details for each experiment, with N per genotype group ranging from 6-16 samples with no obvious outliers and clear separation between nuclear and synaptosomal fractions by PCA (**Figure S5A-C**). We used edgeR^62^ including sequencing batch and sex as covariates. We also analyzed male and female samples separately, finding generally shared DE signatures (similar as for snRNA-seq) (**Figure S5D, Tables S12-15**). Across the four conditions, the largest number of DEGs at either FDR < 0.1 or P < 0.05 was in the synaptosomal preparation, followed by the nuclear preparation, then full cortex from *Chd8^+/-^*and lastly *Chd8^+/flox^* (**Figure 4A**). *Chd8* was among the top downregulated DEGs in all but the nuclear mRNA condition, consistent with nonsense-mediated decay of mutant *Chd8* transcripts outside the nucleus (**Figure 4B**). To identify shared and divergent DE signatures across conditions, we compared logFC of DEGs; shared signatures are represented by concordant LogFC direction, condition-specific effects are captured by patterns of change (up or down) in one condition with LogFC around zero in the other condition, and divergent DE signatures are represented by opposite logFC direction. Comparison between *Chd8^+/-^*and *Chd8^+/flox^* logFC for genes with FDR < 0.1 and P-value < 0.05 in *Chd8^+/-^* (**Figure 4C**) captured largely shared signatures, indicating that effects from post-mitotic heterozygous ablation in excitatory neurons accounts for a significant part of DEG transcriptional pathology. DEGs identified in both the nuclear and synaptosomal conditions (FDR < 0.1 and P < 0.05) showed strong concordance with bulk cortex (**Figure 4D**), however there were striking differences and relatively low concordance between nuclear and synaptosomal enriched mRNA fractions (**Figure 4E**).

**Figure 4:**
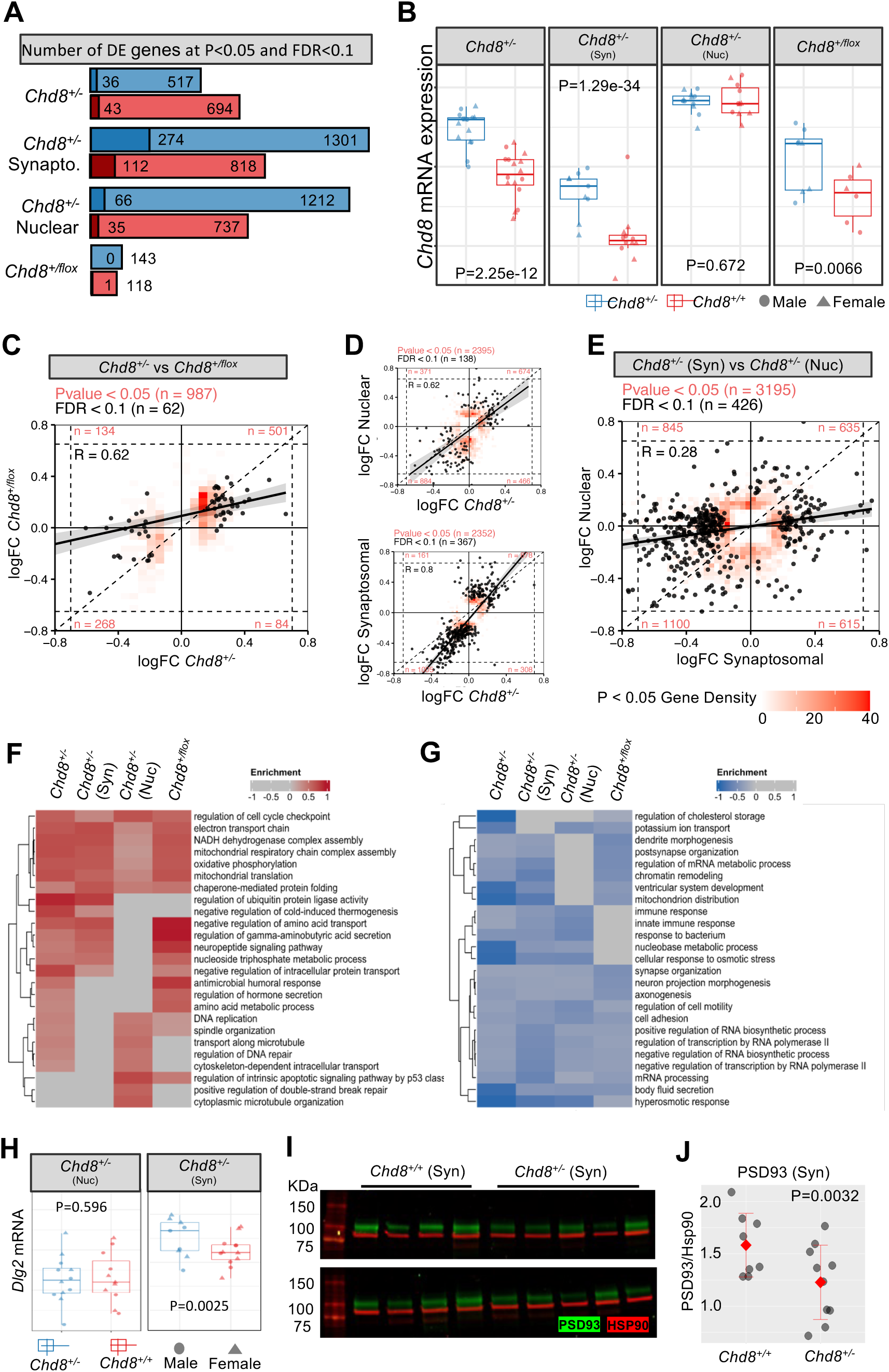
Shared DE in constitutive *Chd8^+/-^* and *Camk2a-CRE* conditional Chd8^+/flox^ mice, with divergent impacts on nuclear and synaptosomal mRNA and protein. A) Barplot of DEG counts at P < 0.05 and FDR < 0.1 across bulk RNA-seq experiments (constitutive *Chd8^+/-^*, *Camk2a-CRE* conditional *Chd8^+/flox^*, and nuclear and synaptosomal preparations from the *Chd8^+/-^* line). B) *Chd8* mRNA is down-regulated in all but the nuclear preparation (P-values from EdgeR). C-E) Concordance plots comparing bulk RNA-seq logFC of DEGs detected in either dataset. Number of DEGs at different criteria listed above, R value for best fit line for FDR < 0.1 is listed on the plot. For P < 0.05, local density is plotted instead of individual data points due to large number of genes. For FDR < 0.1, individual datapoints are shown. The count of P < 0.05 DEGs in each quadrant listed in red. Higher data points in the upper right and lower left indicate logFC directional concordance between experiments. C) Concordance between constitutive and conditional datasets. D) Concordance for nuclear (top) and synaptosomal (bottom) versus whole cortex. E) Reduced concordance between nuclear and synapsomal datasets. F-G) Selected enriched GO terms for upregulated F) and downregulated G) genes across experiments. H) Downregulation of Dlg2 in synaptosomal RNA. I-J) Western blot (I) validating decreased synaptosomal PSD93 protein (J) in *Chd8^+/-^* cortex (N = 8 *Chd8^+/+^*, 10 *Chd8^+/-^*, P-value from Student’s t-test).

To compare pathway-level dysregulation across bulk RNA conditions, we used GSEA to identify Gene Ontology (GO) terms where the annotated gene set exhibited significant shift in LogFC distribution towards up- or downregulation, with shared and condition-specific enrichment for representative terms shown in **Figure 4F-G**. Most of the up- and down-regulated terms identified in the constitutive *Chd8^+/-^* were shared in the conditional *Chd8^+/flox^*, and most terms in the nuclear or synaptosomal enriched conditions were shared in the full cortex preparation, findings that consistent with DEG concordance across contrasts above. We identified shared upregulation of terms including *Cell cycle checkpoint*, *Electron transport chain*, *Mitochondrial translation*, and *Chaperone mediated protein folding*, showing overlap with upregulatd gene modules and DEGs terms that were also detected in glutamatergic neuron subtypes in the snRNA-seq data (**Figure 3**). Among upregulated terms that differed between synaptosomal and nuclear conditions were *Regulation of gamma-aminobutyric acid secretion* and *Neuropeptide signaling pathway* in the synaptosomal and *Regulation of DNA repair* and *Transport along microtubule* in the nuclear fraction. For downregulated terms, *Axonogenesis*, *Synapse organization*, *Cell adhesion*, and *Regulation of transcription by RNA Pol II* were shared, while *Dendrite morphogenesis* and *Postsynapse organization* were example synaptosomal-enriched terms. Bulk RNA-seq results from the full cortex and from the nuclear and synaptosomal conditions showed evidence of correlation with DEG findings from snRNA-seq, though with substantial overall differences in DE signatures (**Figure S5E**). As expected, the nuclear enriched mRNA showed the highest correlation to snRNA-seq nuclear DE, as well as upregulation of genes associated with the glutamatergic M1 and M7 hdWGCNA modules that were broadly upregulated across glutamatergic subtypes in the snRNA-seq data, validating relevance of these broad DEMs (**Figure S5F**). GO terms associated with L2/3 IT enriched upregulated M4 and M9 modules associated with mitochondrial energetics were among the strongest bulk RNA-seq upregulated GSEA hits across conditions. We performed qPCR on 13 DE targets from the constitutive cortex, all 13 genes tested showed the same direction of change as RNA-seq DE analysis (**Figure S5G**). Finally, for the ASD and neuropsychiatric-associated synaptosomal-specific downregulated *Dlg2* gene^63–66^, which encodes the PSD93 excitatory synaptic scaffold protein, we validated reduction at the protein level via Western blot (P = 0.0032, Student’s t-test) in synaptosomal preparations (**Figure 4F-4H**). These results solidify transcriptional pathology in adult cortex due to *Chd8* haploinsufficiency, provide insights into cell-autonomous effects in post-mitotic excitatory neurons, and suggest differential mRNA levels are driven by both transcriptional and post-transcriptional mechanisms that have heightened impact on synaptosomal mRNA levels.

### Altered excitatory neuron spine dynamics and calcium signaling coherence in Chd8^+/-^ mice

The snRNA-seq and bulk RNA-seq both pointed to cortical excitatory neuron dysfunction and previous studies have identified electrophysiological signatures in vivo and in vitro in glutamatergic neurons^21, 36, 37, 40, 41^. To further investigate structural pathology of cortical neurons, we crossed *Chd8^+/-^* mice with thy1-YFP mice, which express cytoplasmic yellow fluorescent protein in a sparse subset of layer 5 pyramidal neurons (L5 PyrNs)^67^, and performed in vivo two-photon (2P) imaging on apical dendrites of L5 PyrNs. We compared the density and dynamics of dendritic spines (postsynaptic structure of majority excitatory synapse) between WT and *Chd8^+/-^* littermates at adolescence (P30) and adulthood (P90) mice (**Figure 5A**). Adolescent *Chd8^+/-^*mice had lower spine density on the apical dendritic tufts of L5 PyrNs (**Figure 5B**). Following the same dendritic segments over seven days, we also found that *Chd8^+/-^*mice had increased dendritic spine formation relative to WT mice, but comparable spine elimination (**Figure 5C**). On the other side, adult *Chd8^+/-^*mice had normal spine density and dynamics (**Figure 5D-E**). These results indicate that *Chd8* haploinsufficiency delays spinogenesis of cortical neurons during early development (before P30).

**Figure 5:**
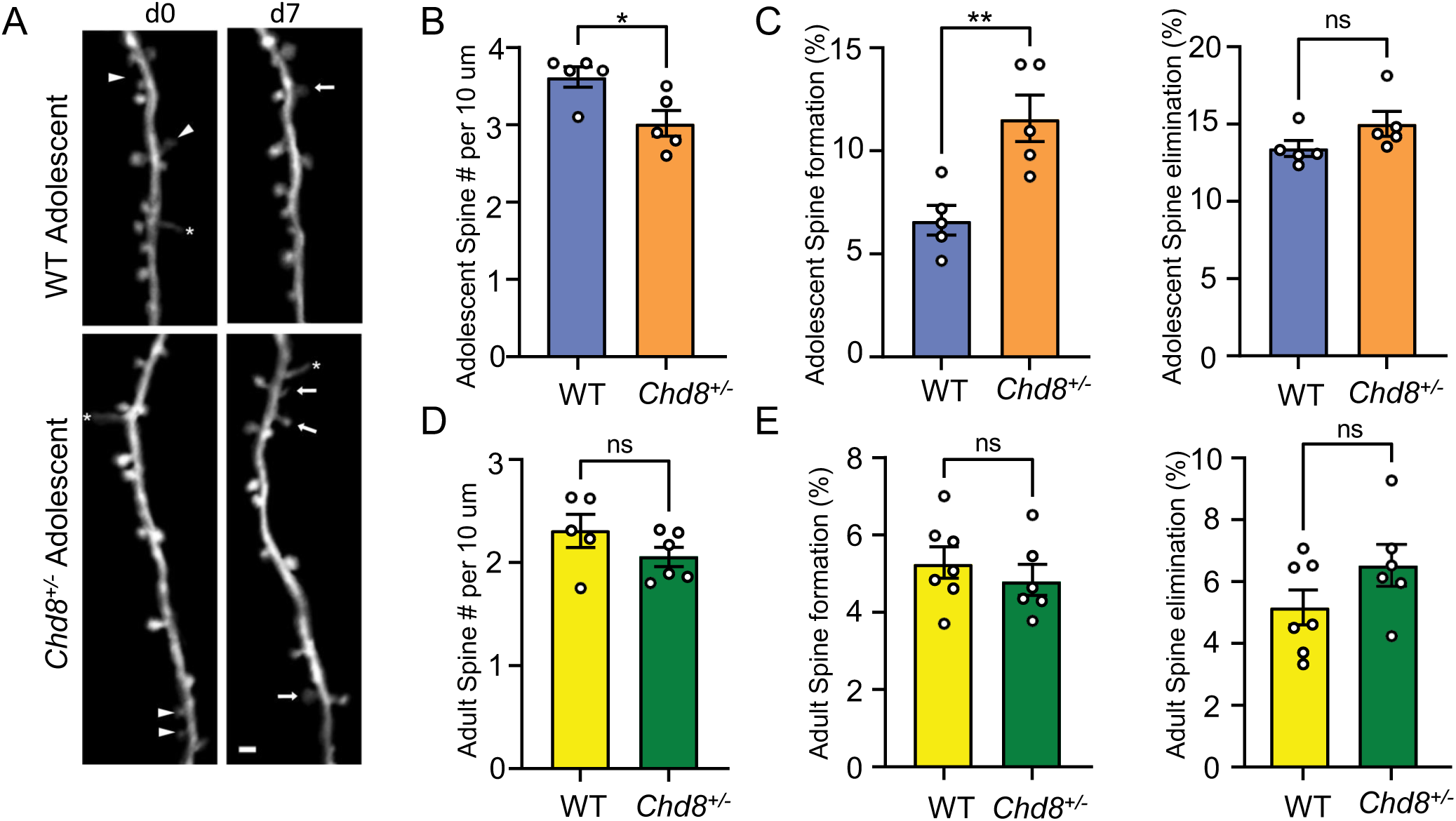
Delayed spinogenesis during *Chd8^+/-^* development. A) Dendritic segment imaged over 7- days in somatosensory cortex of adolescent WT and *Chd8^+/-^*mice. Arrowheads: eliminated spines; arrows: formed spines; asterisks: filopodia. Scale bar = 2 µm. B, D) Spine density in adolescent (b) and adult (D) *Chd8^+/-^* or WT littermates. C, E) The percentage of spines formed or eliminated over 7d is higher in adolescent (D) and adult (E) *Chd8^+/-^*than WT littermates, mean, SEM., unpaired t-test. **p<0.01.

To understand how disruption of *Chd8* signaling affects the function of a cortical microcircuit, we performed calcium imaging in acute slices taken from wildtype or *Chd8^+/-^* mouse brains expressing the genetically encoded calcium indicator *GCamp6f* in prefrontal cortex neurons under the control of the human *Synapsin* promoter (**Figure 6A-D**). We collected 60-minute recordings from a total of 544 neurons from 8 wildtype slices (68+/-9 neurons/slice) and 818 neurons from 10 *Chd8^+/-^*slices (82+/-9 neurons/slice). Overall, the activity of wildtype and *Chd8^+/-^*slices was similar (**Figure 6E-F**; mean fraction of frames active = 5.2+/-0.2% WT vs 5.0+/-0.2% HET, p = 0.4, t-test). To examine the functional connectivity across the prefrontal microcircuit, we binned neurons into eight 75 µm bins starting at the cortical surface and extending to a depth of 600 µm, and calculated the correlation coefficient between the activity of neurons according to the location of each neurons within the slice, then averaged the correlation coefficient based on the depth of 600 µm, and calculated the correlation coefficient between the activity of neurons according to the location of each neurons within the slice, then averaged the correlation coefficient based on the position of each neuron in the cell pair (**Figure 6G-H**). Visual examination of the resulting matrix of averaged correlations revealed that neuron pairs from heterozygous *Chd8^+/-^* slices had on average lower correlation coefficients compared to neuron pairs from WT slices, and this was particularly pronounced when at least one neuron was located in the superficial (< 300 µm from the cortical surface) aspect of the slice (**Figure 6I-L**; mean correlation coefficient WT 0.0057 +/-0.0001 vs 0.0030+/-0.0005, p < 0.05, t-test; n = 8 WT and 10 HET slices; mean correlation coefficient superficial WT 0.0050+/-0.0009 vs HET 0.0021+/-0.0004, p < 0.01, t-test; n = 8 WT and 10 HET slices), suggesting that these neurons are relatively disconnected from deeper layer neurons in the prefrontal microcircuit of mutant *Chd8^+/-^* animals.

**Figure 6:**
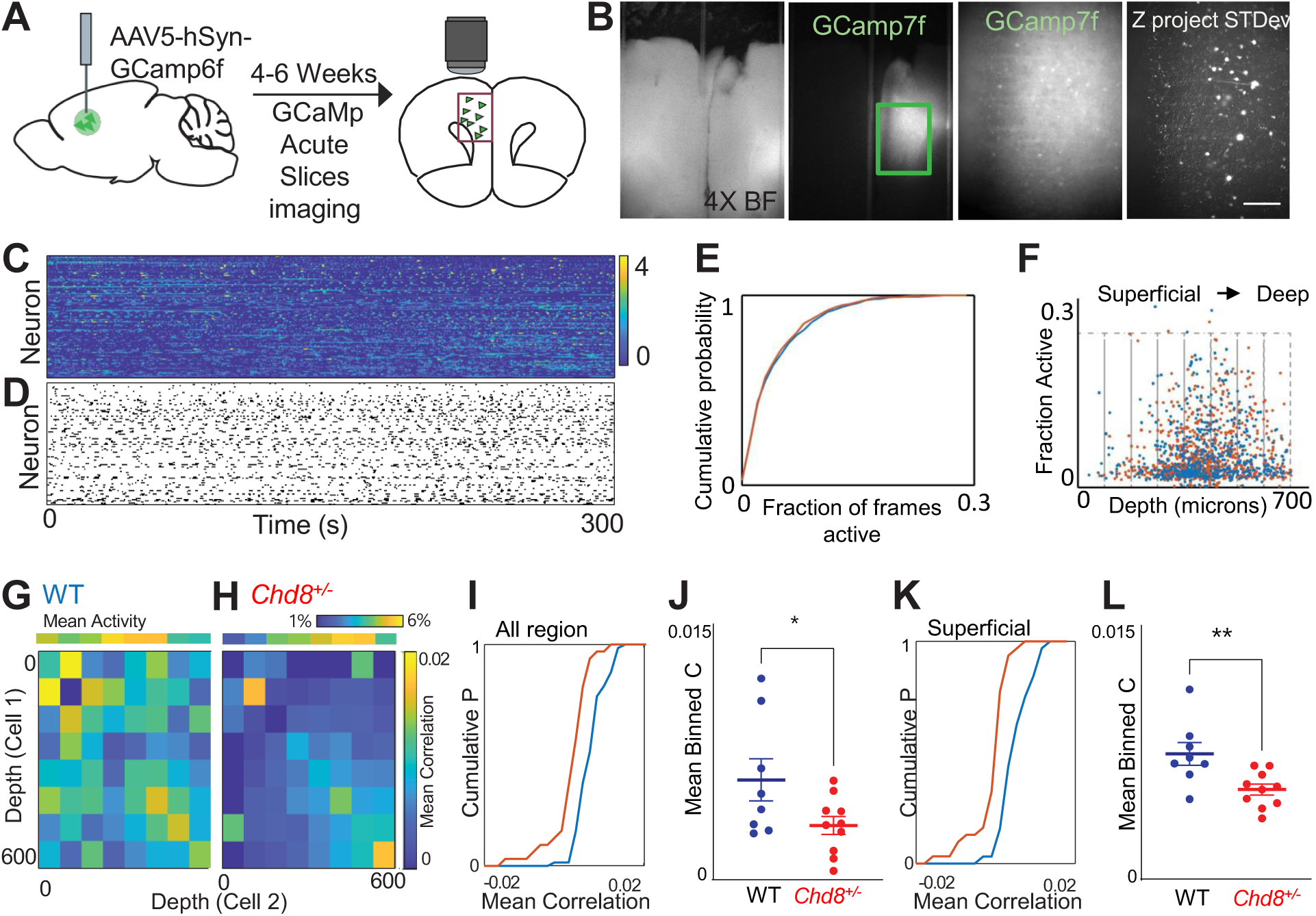
Disruption of coordinated activity in *Chd8^+/-^* PFC microcircuits. A. Mice injected with *Syn-GCamp6f* and activity recorded from acute slices after 4-6 weeks. B, wide field and GFP at 4x (left). 10X of individual neurons and z-projection of standard deviation across frames (right). C, z-scored fluorescence traces from neurons in a single slice. D, Raster of events. E, Cumulative probability function showing overlap in fraction of frames active for neurons from WT (blue) and *Chd8^+/-^*(red) slices. F, Activity of each cell as a function of depth from cortical surface. Cells binned by distance from surface (gray boxes) and mean correlations within and between bins calculated. G, H, 8x8 matrix of mean correlation coefficient calculated for cells within each pair of bins, averaged for WT (G) or *Chd8^+/-^* (H) slices. I, cumulative distribution function (CDF) of mean correlation across 544 pooled WT (blue) neurons and 818 pooled *Chd8^+/-^* (red) neurons for all bins. J, Mean correlation coefficient of WT and *Chd8^+/-^* slices (p < 0.05, t-test). K, CDF of the mean value of the subset of bins in which neuron 1 was < 300 microns from cortical surface. L, Mean correlation coefficient of superficial neurons in WT and HET slices (p < 0.01, t-test).

### Chd8^+/-^ mice exhibit disrupted sleep

Sleep issues are a common symptom in humans with *CHD8* mutations and the transcriptional dysregulation in *Chd8^+/-^* mice featured upregulation of the circadian-associated M7 glutamatergic module. Thus, we used EEG/EMG to continuously record neural activity during a 24-hour period, assigning states of REM, NREM, and Wake, analyzing 11 *Chd8^+/-^*(female=7, male=4) and 8 WT littermates (female=4, male=4) (**Figure 7A-B’**). The total time in seconds that each mouse spent in each state was calculated for each hour (**Figure 7C-D**). For comparison of sleep patterns, total and standard deviation (SD) were calculated by subject using hourly values and compared for light (Zeitgeber Time ZT0-ZT12) and dark (ZT12-ZT24) phases (**Figure 7E-F**). *Chd8^+/-^* mice had a significant increase in time spent in REM in the dark stage (P = 0.036, unpaired t-test with Welch’s correction), and significantly less time spent in NREM (P = 0.008, unpaired t-test with Welch’s correction) and increased time spent in wake state (P = 0.040, unpaired t-test with Welch’s correction) in the light phase. Comparing SD of the time spent per state across hours, *Chd8^+/-^* mice showed higher variability in all states. SD of REM time was significantly higher during both light and dark phases (P = 0.012 and 0.003, respectively), and SD of NREM and Wake time were significantly higher during dark phase (P-values = 0.017 and 0.006, respectively) and marginally increased during the light phase (P = 0.115 and 0.059 respectively). Together results revealed disrupted sleep patterns in *Chd8^+/-^* mice, including more time in REM and time spent in Wake in the light stage, and higher variability in time spent in specific sleep/wake states across the light and dark stages.

**Figure 7:**
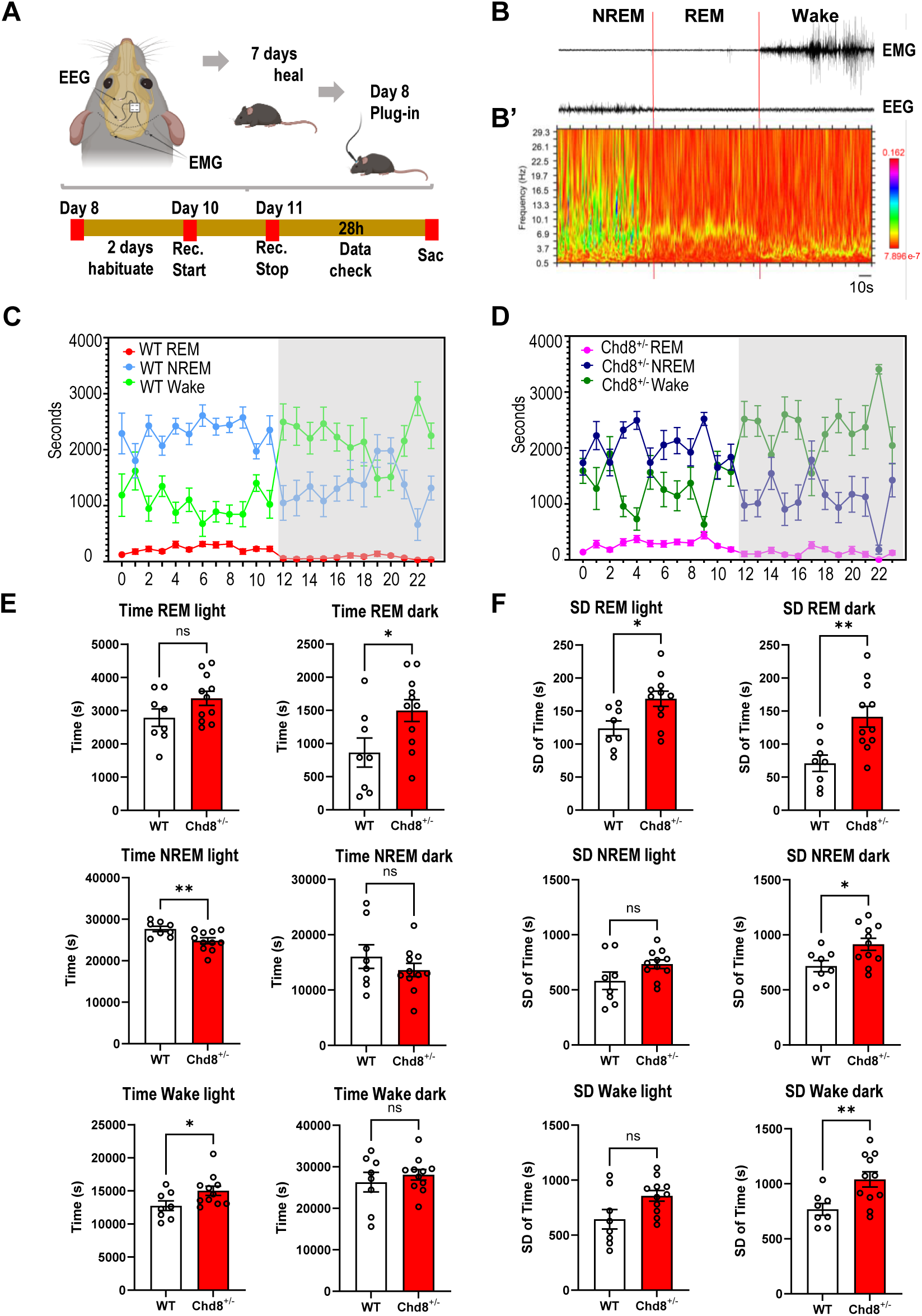
Perturbed sleep patterns in *Chd8^+/-^* mice. A) Schematic of 24hr EEG/EMG sleep analysis. B) Example EEG/EMG traces showing activity during REM, NREM, and Wake state. B’) Spectrogram of the EEG signal in B C-D) Summary of time spent in each state during each hour of the 24hr monitoring period for wild type (C) and *Chd8^+/-^* (D) groups. E) Summary of mean total time spent in each state separated by light and dark stages. F) Summary of SD of time spent in each state. Mean and SD shown, significance form unpaired t-test with Welch’s correction. See text for details.

## Discussion

This study extends understanding of how *Chd8* haploinsufficiency alters the adult mouse cerebral cortex across transcriptomic and functional modalities. Via stereology and snRNA-seq, we show that increased cortical volume in *Chd8^+/-^* mice is not due to increase in cortical excitatory neuron numbers, and, in fact, is associated instead with decreased neuron density. While macrocephaly was not due to increased numbers in excitatory neurons, these cells do exhibit sensitivity to *Chd8* haploinsufficiency at the level of cell-type specific transcriptional pathology, synaptosomal RNA homeostasis, dendritic spine formation, and signaling coherence. Finally, we expand the phenotypic testing of heterozygous *Chd8* mouse models in the domain of sleep, finding perturbations to sleep state that aligns with high prevalence of sleep disturbance reported in patients carrying *CHD8* mutations. Overall, our findings add to the understanding of the effects of heterozygous *Chd8* mutation on the mouse cortex, providing new insights into anatomical, cellular, and circuit impacts, dosage sensitivity during brain development and in post-natal postmitotic neurons, and demonstrating presence of sleep disturbance.

Increased brain size during development and increased cortical volume in adult mice has been reported across different mutant *Chd8* mouse lines^20, 25, 27, 29^. *Chd8* haploinsufficiency has also been linked to altered neurogenesis in mouse^20, 25, 28^ and in vitro in human cellular and organoid brain models^32, 38^. While increased production of cortical excitatory neurons during development leading to increased neurons in the adult cortex seemed a plausible explanation for cortical volume increase, results from both stereology and snRNA-seq show this to not be the case in *Chd8^+/5bpdel^*mice. In fact, we found the opposite pattern of decreased excitatory neuron density driven by increased volume without increased neuron counts or increased soma size. There was a trend of decrease in proportions of certain glutamatergic types in snRNA-seq data, reaching significance for decreased in L6 IT neurons in the mutants. snRNA-seq indicated significant increase in proportion of OPCs and trend towards increase in astrocytes, raising the possibility that increased proportions of non-neuronal cells may explain some of the increased cortical volume. Whatever the roots of *Chd8*-associated macrocephaly, it is unclear what the link between this neuroanatomical signature and relevant social and cognitive phenotypes, with a recent study finding little correlation between adult brain weight and behavioral phenotypes for *Chd8^+/-^* mice from a series of genetic backgrounds^34^. It could be that causes of macrocephaly and behavioral phenotypes in *Chd8^+/-^*mice are independent, or that the mechanisms for are linked, but with variable anatomical and behavioral outcomes in adult mice. Our results do point to glutamatergic pathology and one intriguing possibility is that changes in neuronal morphology and microstructure organization, for example dendritic arborization, could account for the changes. Further studies will be needed to resolve specific causes of volume increase in *Chd8^+/-^* adult cortex.

Via unbiased snRNA-seq, we found that glutamatergic neurons exhibit the strongest transcriptional pathology across cell types in adult mouse *Chd8^+/-^* cortex, with the most extensive impacts in L2/3 IT and L4/5 IT subtypes. We identified subtype-specific and shared glutamatergic DEMs, including shared upregulation of M1 and M7 modules, which are associated with general neuron-associated pathways and with specific circadian rhythm GO terms. Transcriptional pathology in bulk RNA-seq analysis of constitutive and conditional *Chd8^+/-^* mutants here aligns with *CHD8* studies in vitro and using mouse models^12, 20, 22, 24, 25, 28, 68^, with our study adding substantially via integrating single cell and bulk, constitutive and conditional mouse models, and synaptosomal mRNA fractions. These findings include upregulated aerobic respiration and mitochondrial energetics, which we map to having the strongest signature in L2/3 IT neurons, and downregulation associated with synaptic organization and signaling. Bulk RNA-seq transcriptional changes in excitatory neuron-specific *Chd8^+/flox^* mice paralleled changes in constitutive heterozygous *Chd8^+l-^* mutants, demonstrating continued requirement for full *Chd8* dosage and indicating that the majority of excitatory neuron transcriptional phenotypes in mature *Chd8^+/-^*mice may be independent of developmental impacts. Our findings of effects in *Chd8^+/flox^* mice due to ablation in excitatory neurons are consistent with effects reported in earlier in vitro studies of both heterozygous and homozygous neuronal ablation^40, 69^, and extend previous studies of adult homozygous mutant *Chd8^-/-^* conditional mutant mice that did not examine NDD-relevant heterozygous mutants^44^. We further link transcriptional dysregulation in glutamatergic neurons to functional impacts in these cells, finding altered dendrite spine formation during adolescence which manifests at the circuit level as de-correlated spontaneous prefrontal activity. Finally, though not directly linked to specific cells or circuits, we identified perturbed sleep patterns *Chd8^+/-^*. These findings are highly relevant to NDDs, as disrupted synaptic development and altered network activity represent candidate causal mechanisms and sleep disturbances are commonly reported in individuals with ASD and related conditions^70^ and are one of the more penetrant phenotypes in CHD8-NDD^71^. We previously tested *Chd8^+/5bpdel^*mice across a battery of behavioral assays, finding deficits in novel object recognition and fear conditioning in the learning and memory domain with no differences in sociability using the three-chamber social preference test or in male-female interactions^20^. Other constitutive *Chd8^+/-^* mouse studies similarly have not identified classic sociability deficits in the three-chamber social preference assay and behavioral and cognitive phenotypes vary across *Chd8^+/-^* studies and across mouse genetic backgrounds^20–22, 24, 25, 27, 34, 37^. As such, behavioral studies of *Chd8* mutant mice have proved challenging to interpret. Our results, along with studies that have identified other phenotypes at cellular, molecular, and anatomical level such as via electrophysiology or structural MRI, highlight molecular endophenotypes that may be more consistent in expression and have high translational relevance. Future studies will be needed to determine which aspects of *Chd8^+/-^*mouse phenotypes are driven by developmental versus later requirement for full dosage, which specific neuronal and glial populations are dependent on *Chd8*, and if molecular, cellular, and neuroanatomical phenotypes are consistent between mutant lines and across mouse and human *CHD8^+/-^* models.

One novel finding from our study was the significant differences in DE signatures between nuclear and synaptosomal mRNA preparations. Bulk nuclear RNA-seq recapitulated upregulation observed for the two glutamatergic DEMs with broad dysregulation and had the highest concordance with snRNA-seq DEGs, as well as being strongly concordant with the full bulk cortex RNA-seq. Thus, nuclear RNA changes observed at both single cell/nucleus and bulk level appear robust. Surprisingly, there was relatively weak concordance between synaptosomal and nuclear DEGs, with a stronger overall DE signature in the synaptosomal enriched preparations. These findings indicate disconnect between changes in transcriptional rate, captured by nuclear RNA datasets, and changes in steady-state mRNA levels at the synapse in *Chd8^+/-^*cortex. We and others previously identified altered mRNA splicing in *Chd8^+/-^*mouse and *in-vitro* models^20, 72^, raising the possibility that *Chd8* haploinsufficiency directly or indirectly alters RNA processing. ChIP-seq studies of CHD8 have consistently identified enrichment of regulatory targets associated transcription and RNA processing^73^. We did observe down-regulation of RNA processing associated with L4/5 IT specific DEM M6, though this was specific to this population. Beyond a better understanding of the molecular function of Chd8, future studies are needed to determine whether the synaptosomal DEG signature is specific to synaptosomal localized mRNA, or a more broadly present in cytosolic mRNA overall, what the cell-type specific impacts are on mRNA processing and localization, whether synaptosomal RNA differences are associated with differences in synaptic protein makeup and synaptic function, and if this finding extends to *CHD8^+/-^* human neurons. Of more broad relevance, our findings of altered post-transcriptional RNA dynamics link CHD8 with other ASD/NDD genes with primary function in RNA processing, for example *FMR1*/FMRP^74, 75^, and our finding of pathology arising do to disruption of this function may by a more broad hallmark of ASD^76, 77^.

There are limitations to this work that should be considered. The number of replicates (N=5 per genotype) was cost-limited for snRNA-seq, somewhat limiting statistical power. Additionally, nuclei-based methods may miss changes in cytosolic mRNA levels^78, 79^, which may be relevant based on our bulk analysis. Further, there are challenges with snRNA-seq for DE testing at single gene level using individual cells/nuclei. While Wilcoxon tests are commonly used in single cell differential expression analysis, we found this method yielded high false positive rates (as issue that has been raised previously^80^). In contrast, pseudobulk and hdWGCNA methods take advantage of aggregating cells or genes to compare expression at the cluster or module level, giving more robust DE results at the cost of reducing resolution for single cell and single gene level effects. Finally, it is unclear how much the transcriptomic pathology extends to protein level or impacts function. With regard to functional studies, we chose three *in vivo* assays to focus on: spine dynamics, correlated GCaMP signaling, and sleep. Previous studies have not tested these phenotypes, and we believe our results provide new insights into pathology. However, other assays would be valuable and each of the domains we studied could be investigated at higher depth. We did not test conditional *Chd8^+/flox^*mice beyond bulk RNA-seq, leaving unanswered whether this model recapitulates other pathology. Finally, *Chd8* mutant mouse lines exhibit variation in phenotype penetrance and expressivity, which is at least in part due to the impact of genetic background^34^. Considering this, our findings may not be conserved in other *Chd8* mouse models, though we note overlapping transcriptional signatures described here and in other studies as summarized above. While these limitations hold, our results lay the foundation for future work characterizing constitutive and conditional mutant *Chd8* mouse models.

In summary, our study extends understanding of the impacts of *Chd8* haploinsufficiency in vivo in mouse brain, with a focus on glutamatergic excitatory neurons. We show that constitutive and conditional heterozygous ablation leads to convergent transcriptional pathology and to impacts on neuronal spinogenesis and circuit function. Our findings illustrate how loss of *Chd8*, which has received attention for high expression in early development and role as a chromatin remodeler, impacts the cell-type specific makeup and pathology of the adult cortex and impacts post-transcriptional mRNA dynamics, extending understanding across molecular, cellular, circuit, and organismal phenotypes. Our findings support a framework where excitatory neuron dysfunction at the level of cortical excitatory synapses and circuits may be an area of casual convergence in NDDs and raising the possibility that interventions targeting postnatal postmitotic cortical neurons may be effective at addressing at least some of the underlying NDD pathology.

## Acknowledgements

This work was supported by the National Institutes of Health National Institute of Mental Health (NIH NIMH) R01 MH120513 to A.S.N., National Institute of Neurological Disorders and Stroke (K08 NS105938-01A1 to N.A.F.), the Simons Foundation (Award # 604325) and the National Institute of Mental Health (R01MH117961 to V.S.), as well as the UCSF Dolby Family Center for Mood Disorders. C.P.C. was supported by the NIH NIMH Autism Research Training Program fellowship (T32 MH073124-16). S.L. was supported by T32 GM 007377 and F31 HD113328-01A1. A.A.W. was supported by NIH NIMH F31 MH119789 and T32 GM007377. We thank Dr. Wenjie Bian and Dr. Luis de Lecea at Stanford University, for their technical advice on EEG/EMG experiments and data analysis. We thank Dr. Caitlin Moyer for her technical assistance and for contributing to the execution of the spine imaging analysis.

## Author contributions

A.S.N. conceived the project with substantial contributions from Y.Z., D.A. and V.S. C.P.C. led the project execution, designed experimental work and oversaw most aspects of data generation and analysis. S.A.L. fully executed snRNA-seq bioinformatics and led most of the data analysis with contributions from K.C., N.S., E.F., N.C., E.S., C.A., A.W. and A.S.N. C.P.C., W.A., J.B., and S.F. carried out experimental stereology and analysis. C.P.C., S.A.L., P.B., A.K.S., D.P., D.R., S.M., M.C., L.D., J.Z. and R.O. carried out experiments. C.M. and J.B. performed 2P spine imaging and analysis, J.S., B.M., and J.L. performed EEG recording and analysis. C.P.C., S.A.L. and A.S.N. wrote the manuscript.

## Methods

### Animals

Generation of Cas9-mediated 5bp frameshift deletion in exon 5 of *Chd8* in mice (here referred to as *Chd8^+/-^* mice) was previously described^20^. For in vivo imaging, this mouse line was crossed with the thy1-YFP-H (JAX strain: 003782) mouse line and maintained in the C57BL6/J (JAX strain: 000664) background. Mutant mice *Camk2a-CRE Chd8^+/flox^* were obtained by crossing Chd8 loxP mice on C57BL/6J (*Chd8^em3Lutzy^*; JAX strain: 031555) to Camk2a-CRE (*B6.Cg-Tg(Camk2a-cre)T29-1Stl/J*; JAX strain: 005359) to achieve heterozygous ablation in post-mitotic forebrain glutamatergic neurons. All protocols utilized in the generation of mouse brain samples were approved by the Institutional Animal Care and Use Committees (IACUC) at the University of California Davis and University of California Santa Cruz. Mice were housed in a temperature-controlled vivarium maintained on a 12-h light–dark cycle. We made efforts to minimize pain, distress and the number of animals used in the study.

### Histological tissue preparation and Nissl Stain

Thirty-six 42-day-old mice were included in this analysis (16 brains per genotype, 16 per sex). Mice were anesthetized using Isoflurane and perfused transcardially using room temperature saline solution (0.9% NaCl) followed by 4% paraformaldehyde (PFA) in 0.1 M phosphate buffered (PB) saline using a peristaltic pump (Minipuls 3, Gilson) at 20 rpm flow rate. Brains were removed and post-fixed in 4% PFA at 4°C for 24 h. Fixed brains were cryoprotected in 50 ml of 10% and 20% glycerol solutions with 2% DMSO in 0.1 M phosphate buffer (PB; pH 7.4; for 24 and 72 hours, respectively). Brains were blocked at -2.00mm posterior to the interaural line prior to flash freezing. Sections were cut using a freezing, sliding microtome in 4 series at 60 µm (Microm HM 440E, Microm International, Germany). Every 4th section was collected in 10% formaldehyde in 0.1 M PB (pH 7.4) and post-fixed at 4°C for 1 week^81^. Sections were rinsed twice for 1 hour each in 1ml of 0.1M PB, mounted on gelatin-coated slides, and air-dried overnight at 37°C. Defatting was performed in a mixture of chloroform/ethanol (1:1, vol) for 2 x 1 hour. The brains were rehydrated in graded alcohols; 100% ethanol, 100% ethanol, 95% ethanol, 70%, 50% and dipped in deionized water^81, 82^. Sections were stained 40 seconds in a 0.25% thionin solution (Fisher Scientific, Waltham, MA, cat. no. T-409), dehydrated through graded alcohols, differentiated in 95% ethanol and 5 drops of glacial acetic acid, cleared in xylene and cover-slipped with DPX (BDH Laboratories, Poole, UK)^81^.

### Stereology

The inferior boundary of the cortex was defined as the bottom of the agranular insular area (AIA), and the superior boundary was defined as the anterior cingulate area (ACA) or the retrosplenial area (RSA) and the white matter was then followed to the ACA/RSA border^83^. Neuron number, neuronal soma size, and brain volume were estimated using Stereoinvestigator V10.50 (MBF Biosciences, Williston, VT) attached to a Zeiss Imager.Z2 Vario with a Zeiss AxioCam MRc. An EC PlanNeoFluar 2.5 x objective was used to trace the outline of the cortex and a PlanApoChromat 100x oil objective was used for cell counts and soma size determinations. The volume of the cortex was determined using the Cavalieri method^53, 54, 84^. For each mouse, 15-18 sections (480 µm apart) were analyzed with the first section chosen randomly amongst the first through the cortex. Estimates of neuron numbers were obtained using the Optical Fractionator method^55, 85^. Since no lateralization was found in the volumetric estimates (p = 0.65), we sampled the left hemisphere of half of the mice and the right hemisphere of the other half for estimates of neuron numbers. We used the same sections that were used for volumetric estimates. We used a counting frame size of 18x18x10µm with 2 µm guard zones, and a scan grid of 480x480 µm placed at a random angle. Neurons were defined following established protocols^53–55, 84–86^ based on: 1) dark staining, 2) a well-defined nucleolus and 3) a round, non-jagged cell body^81, 86–89^. Section thickness was measured at every counting site. The volume of neuronal somas was determined using the Nucleator method^90^ during the Optical Fractionator analysis. Comparison of stereological measures (volume, neuron number, and soma size) were performed via ANOVA to examine global effects of sex and genotype and test for sex by genotype interaction, and then via t-test to compare the effect of genotype in males and females. In addition to test results, effect sizes are reported for ANOVA (partial eta squared, *ηp*^2^) and t-test (Cohen’s d). Statistical analyses were performed using SPSS (version 26), plots were generated using *ggplots2* package in R programming language^91^.

### Rostro-Caudal Volume analysis

To investigate the effects of genotype and sex on Cavalieri counts, mixed-effects regression models were employed using the ‘lme4’ and ‘lmerTest’ packages in R. The initial model tested the main effect of genotype on Cavalieri counts while accounting for sex, batch effects, and the z-position of the mouse brain sections. Polynomial terms up to the fourth degree for z-position (Zpos) were included to capture potential non-linear relationships observed in exploratory data analysis. Subsequently, a model was fitted to examine the interaction between genotype and sex, including an interaction term for these variables. This model was structured to evaluate whether the effect of genotype on Cavalieri counts varied by sex. The analysis was extended to assess the effects of genotype across different rostral-caudal brain segments. The segments - Rostral, Median, and Caudal - were defined based on changes in direction observed in the z-position plot, aiming to reflect the anatomical structure of the mouse brain. Dummy coding was applied to compare each segment against the reference levels. The final model included these segments as factors, alongside the previously mentioned covariates and polynomial terms for z-position. For data visualization, the ‘ggplot2’ package was utilized to create plots depicting the relationship between z-position and Cavalieri counts. Separate smoothed plot lines were generated for each sex and genotype combination, providing a clear visual representation of the data across different brain regions. All statistical analyses were performed using R. The mixed-effects models allowed for robust estimation of fixed effects while accounting for random effects associated with batch and individual mouse variations.

### Adult mouse cortex dissections

Mice were anesthetized with isoflurane and euthanized via cervical dislocation in accordance with institutional animal care protocols. Brains were rapidly extracted, and cerebral cortices were dissected on ice following standard anatomical landmarks. Dissected tissues were snap-frozen in liquid nitrogen or dry ice to preserve molecular integrity and minimize RNA degradation, and stored at -80°C until further use. These samples were subsequently used for RNA extraction, nuclear isolation, or cellular fractionation, as described below.

### Cellular Fractionation (synaptosomal preparation)

Brains from *Chd8^+/-^* and wild-type littermates were collected on or around PND60. Following cervical dislocation, brains were sub-dissected to isolate Cortex. Flash frozen cortices were stored at -80C until further processing. On the day of cellular fractionation, samples were treated with Syn-Per reagent (ThermoFisher Scientific Catalog #87793) to obtain cytosol, synaptosome and flowthrough (“total”) fractions. The “total” fraction was then used treated with “NE-PER™ Nuclear and Cytoplasmic Extraction Reagents” (cat #78833) to obtain the nuclear, and a “cleaner cytosol” fractions. Total RNA from these fractions was then extracted using RNA RNAqueous™ Total RNA Isolation Kit (cat# AM1912 ThermoFisher Scientifics). Manufacturer protocols were followed with minor adjustments to increase RNA yield from cellular fractions. Adjustments included combining the cortex samples with 500μl of Syn-Per reagent and 5μl of protease inhibitors during homogenization, based on our starting sample amount, and adding 350μl of RNA extraction lysis buffer immediately after fractionation to the synaptosome and nuclear fractions.

### RNA Isolation, bulk RNA-Sequencing and bioinformatics analysis

Mouse cortices were collected at around PND60 from the following groups: 15 wild-type (WT) mice and 16 *Chd8^+/-^* mice (8 males and 7 females in the WT group; 7 males and 9 females in the *Chd8^+/-^* group), as well as 8 WT and 6 *Chd8*^+/flox^ mice (5 males and 3 females in the WT group; 4 males and 2 females in the *Chd8^+/flox^* group). Samples were snap frozen on dry ice and stored at -80C until RNA preparation. Total RNA was obtained using RNAqueous Total RNA Isolation Kit (cat# AM1912 ThermoFisher Scientifics), and assayed via Agilent RNA 6000 Nano Bioanalyzer kit/instrument. Sample RIN scores ranged from 8 to 9.4. Poly-A-enriched mRNA libraries were prepared at Novogene using Illumina reagents. Libraries were sequenced using Illumina NovaSeq 6000 S4 system, paired-end 150 (PE150) method. Reads were aligned to mouse genome (GRCm38/mm10) using STAR (version 2.5.4b)^92^, and gene counts were produced using featureCounts^93^. On average, 50 million paired-end reads (PE) aligned per sample, with a range of 32 to 84 million reads. Data quality was assessed using FastQC^94^, and principal component analysis (PCA) was used to determine presence of sample outliers. All 24 samples were qualified for the analysis. Raw RNA-seq fastq files and a gene count matrix is available on GEO (GSE300997). Bioinformatic analysis was performed using R programming language version 4.2.1 (R Development Core Team, 2015) and RStudio integrated development environment version 2023.06.0 (Team R, 2018). Plots were generated using ggplot2 R package version 3.4.0. Heatmaps were generated using pheatmap R package 1.0.12. In addition to total cortical RNA-seq, bulk RNA-sequencing was also performed on nuclear and synaptic fractions isolated from the cortex of *Chd8^+/-^* (germline heterozygous) and wild-type littermates at PND60. A total of 13 *Chd8^+/–^*and 12 wild-type nuclear fraction libraries were sequenced from both male and female mice. Following quality control (**Figure S5C**), 9 mutant (4 male, 5 female) and 12 wild-type (7 male, 5 female) nuclear libraries were retained for downstream analysis. For the synaptic fraction, 11 wild-type (6 male, 5 female) and 9 mutant (3 male, 4 female) libraries were sequenced, of which 7 mutant and 11 wild-type libraries passed QC and were used for downstream analysis. RNA from both fractions was processed using the same RNA isolation and sequencing protocols as described above. Raw and processed data for these samples are also available through GEO (GSE300997).

### Bulk differential expression (DE) analysis

For DE analysis we used edgeR R package^62^. Genes with a minimum of 1 counts per million (CPM) in at least six samples were included in the analysis. The first surrogate variable from the SVA-batch-correction method was used as a covariate in the edgeR GLM model^95^. For sex-stratified DE, we used a threshold of CPM > 1 in at least 3 samples, and no batch correction. Reads Per Kilobase per Million mapped reads (RPKM) were used for plotting summary heatmaps and expression data of individual genes.

### Bulk gene ontology enrichment analysis

To test for enrichment of GO terms we used the TopGO R package version 2.34^71^. Mouse Gene Ontology (GO) data was downloaded from Bioconductor (org.Mm.eg.db). For the analysis presented here, we restricted our testing to GO Biological Process annotations and required a minimal node size of 20, and at least 2 significantly DE genes in a GO term. We used the internal ‘weight01’ testing framework and the Fisher exact test, a strategy recommended for gene set analysis that generally accounts for multiple testing comparisons. For GO BP analysis, we reported terms with p-value<0.05. For all enrichment analysis, the test set of DE genes was compared against the background set of genes expressed in our study based on minimum read-count cutoffs described above.

### Quantititavice PCR (qPCR) target validation

qPCR was performed using the Analytik Jena qTower³ thermal cycler and PowerSYBR reagent (Thermo Fisher Scientific), following the manufacturer’s protocols. Primers were custom-designed and ordered from Integrated DNA Technologies or Sigma-Aldrich (listed in Table S16). The thermal cycling conditions were as follows: 10 minutes at 95 °C, followed by 40 cycles of 15 seconds at 95 °C and 60 seconds at 60 °C. Relative RNA expression levels were calculated using the comparative Ct (ΔΔCt) method, normalized to the geometric mean of housekeeping genes. Fold changes were computed using the 2^-ΔΔCt formula as described by Pfaffl^96^, with control samples serving as the calibrator.

### Western blot analysis

Synaptosomal fractions were obtained from cortical extracts from PND 60 micethe using the Syn-Per reagent following manufacturer instructions as described above. Protein concentration was quantified using a BCA protein assay kit (Pierce, 23225) and samples were stored at -80℃ until use. For the SDS gel electrophoresis, samples were diluted in 6X Laemmli SDS buffer (375mm Tris-HCl, 9% SDS, 50% glycerol, 0.03% Bromophenol blue) and 5% β-mercaptoethanol, boiled at 70℃ for 10 min, and separated on a 4-20% polyacrylamide tris-glycine protein gel (BioRad). Prior loading into the gel, samples were incubated at 70℃ for 10 min. After SDS-PAGE, proteins were wet transferred onto a PVDF membrane (Millipore Sigma) overnight at 4℃ (13mA, constant current). Membranes were blocked with Intercept PBS blocking buffer (Li-Cor) at room temperature for one hour. Primary antibody PSD93 (1:50, DSHB, Clone N18/28, supernatant fraction) was diluted in 7.5 ml Intercept PBS blocking buffer with 0.1% Tween. Membranes were incubated with the primary antibody solution overnight at 4℃, then washed four times for 10min with PBS with 0.1% Tween (PBST). Fluorescently tagged secondary antibody (Li-Cor) were diluted in 10ml Intercept PBS blocking buffer with 0.1% Tween. After the initial washes, blots were incubated with the secondary antibody solution for one hour at room temperature. Blots were washed an additional four times for 10min with PBST and two times with PBS. Bands were visualized using the Odyssey DLx imaging system (Li-Cor).

### Single Nucleus RNA-seq

#### Nuclei isolation for snRNA-seq and pooling strategy

Frozen issue from the first 6 samples was processed into nuclei using 2mL glass dounce homogenizers containing 2mL of EZ Prep lysis buffer each. Tissue for 3 samples was dounced over ice, 25 times with pestle A and 35 times with pestle B. Each solution was transferred to a 15 mL conical tube, then the same douncing process was repeated for the second set of 3 samples. Solutions from the latter 3 samples were pooled with those of the first 3 such that each pool contained one male and one female of opposite genotype. An additional 6 mL of EZ Prep lysis buffer was added to each conical tube and the tubes were incubated on ice for 5 minutes. Homogenates were centrifuged at 500xg for 5 minutes at 4°C, the supernatant was discarded, and the pellet was washed twice in NSB (1X PBS with 0.01% BSA and 0.1% RNAse inhibitor). The final pellet was resuspended in 1mL NSB then passed through a 20uM cell strainer. Nuclei were counted using a LUNA-FL cell counter and Acridine Orange/ Propidium Iodide stain. The 10X Genomics nuclei isolation kit, with protocol as described, was used to isolate nuclei for the final 4 samples. These samples were combined into 2 pools, each containing one male and one female of opposite genotype, then nuclei were counted as described previously. Nuclei suspensions had concentrations adjusted to target 10,000 nuclei for sequencing per pool, then used as input for the 10X Genomics 3’ Gene Expression Assay. Libraries were generated according to the v3 protocol by the UC Davis DNA Technologies Core. For the first 3 pools, libraries were sequenced with the Illumina NovaSeq 6000 at 80,000 reads per nucleus. The remaining two pools were sequenced with the Element Biosciences AVITI at 80,000 reads per nucleus. Raw fastq files and integrated Seurat object available on GEO (GSE301511).

### RNA-seq relevant bioinformatics

#### Pre-processing of snRNA-seq data and initial QC

De-multiplexed sequencing data were obtained from the UC Davis DNA Technologies Core as FASTQ files. FASTQ files were passed through the Cell Ranger *count* pipeline for alignment to the mouse mm10 genome, filtering, and creation of feature-barcode matrices. Ambient RNA removal was performed using SoupX (version 1.6.1), and Seurat objects were generated using Seurat (version 4.3.0). These Seurat objects were subjected to initial clustering, and each resulting cluster was filtered based on cluster-specific thresholds for UMI count, feature count, and percentage of mitochondrial reads, which were determined using standard deviation criteria. Mitochondrial genes were then removed from all objects.

#### Doublet removal

The DoubletFinder package (version 2.0.4) was used to assign nuclei as ‘Singlet’ or ‘Doublet’. Predicted doublets were removed before proceeding with downstream analyses.

#### Assigning sex and genotype

Nuclei were assigned as ‘male’ or ‘female’ using a custom-built classification tool (https://github.com/nickolas-chu/sxreveal) based on the expression of sex-linked marker genes located on the X and Y chromosomes. *Chd8* genotype was assigned based on the predicted sex and pool-specific metadata. Because each pool consisted of only one male and one female of opposite genotype, all female nuclei in a pool were assumed to be one genotype and all males were assumed to be the other.

#### Clustering analysis

Seurat objects for each of the 5 samples were individually passed through *SCTransform* to normalize counts, then all 5 were integrated using *IntegrateData*. The ‘RNA’ assay of the integrated object was normalized and scaled using *NormalizeData* and *ScaleData*. Principal Components Analysis was performed on the ‘Integrated’ assay of the integrated data to identify top variable genes, and *FindNeighbors* was used to construct a KNN graph based on their euclidean distance in PCA space. The data were then passed through *FindClusters* to group cells into clusters using the Louvain algorithm. Clusters were visualized with uniform manifold approximation and projection (UMAP).

#### Cell type annotation

*FindAllMarkers* was used to identify cluster marker genes. Modules of cell type marker genes described in published scRNA-seq datasets from adult mouse cortex were constructed. Canonical marker genes were added to these modules, then *AddModuleScore* was used to calculate relative expression of each module in each cluster. The cell type associated with the marker gene module that had the highest expression in each cluster was assigned as that cluster’s identity. All nuclei were assigned identities at 3 levels of classification, with ‘Cell_Type_1’ being the most specific and ‘Cell_Type_3’ the broadest.

#### Cell type re-clustering

Clusters that were poorly clustered, had a high percentage of mitochondrial reads relative to other clusters, or had ambiguous marker genes (i.e. markers for multiple distinct cell types) were removed before subsequent analysis. The filtered Seurat object was then re-normalized, re-scaled, and re-clustered as described previously.

#### Differential expression (DE) analysis

A pseudo-bulk approach was used to identify differentially-expressed genes in each cortical cell population. Each cluster was extracted as a subset of the main Seurat object, then raw counts for protein-coding genes were passed through the *DESeq2* pipeline to compare *Chd8* mutant and wild-type nuclei within that subset. Batch was included as a covariate in the *DESeq2* statistical model. DEGs with an adjusted p-value < 0.05 were deemed significant. This analysis was performed at each of the 3 levels of cell type assignment.

#### Burden analysis

A minimum expression threshold was determined based on the minimum expression level (‘baseMean’) of DEGs with p-value < 0.05. For each cell type, DE burden was calculated as the percentage of DEGs relative to the total expressed genes that passed the minimum expression threshold. A permutation test with 1,000 iterations was performed to assess the statistical significance of the observed DE burden. This analysis was performed at each of the three levels of cell type classification.

#### Cell type proportion analysis

For each cell population, nuclei of each genotype were counted, then counts were converted to a proportion of the total cells in each sample. Linear models adjusted for genotype, sex, and batch were used to assess whether or not the proportions of mutant and wild-type nuclei for each cell type were statistically different.

#### hdWGCNA

High-dimensional weighted gene co-expression network analysis (hdWGCNA) was applied to construct gene co-expression networks for each cell population using the hdWGCNA package (version 0.2.16). Nuclei of specific cell types were grouped by sample and passed through the *MetacellsByGroups* function to create metacells. Custom parameters for ‘k’, ‘max_shared’, and ‘min_cells’ were chosen for each cell type. We determined the optimal soft power using *TestSoftPowers*, built the co-expression network with *ConstructNetwork*, and detected modules via hierarchical clustering. Harmonized module eigengenes (hMEs) were calculated for each module using *ModuleEigengenes* and *GetMEs*.

#### Differential module expression analysis

To test for differences in module gene expression by genotype, we analyzed each module identified in glutamatergic neurons by plotting the log fold change (logFC) of its member genes based on pseudo-bulk differential expression analysis. We then compared the median logFC of each module to that of the corresponding ‘grey’ module using a t-test, applying Bonferroni correction for multiple comparisons. Modules with a Bonferroni-adjusted p-value < 0.05 in at least one glutamatergic subtype were considered significantly differentially expressed.

#### GO enrichment analysis

GO analysis was performed on significant DEGs and DEMs using the *enrichGO* function from the clusterProfiler R package^60^ (version 4.10.0), with the gene universe being set to all genes expressed in the cell type the DEGs or DEMs were identified in. Genes and modules were annotated with biological process and molecular function terms. Terms were deemed significant if they had a corrected p-value < 0.05.

#### Gene set enrichment analysis (GSEA)

Gene set enrichment analysis was performed on DEGs from bulk and pseudobulk DE analysis using clusterProfiler with the org.Mm.eg.db database (version 3.18.0, Carlson M (2022). _org.Mm.eg.db: Genome wide annotation for Mouse_. R package version 3.15.0.). For pseudobulk DEGs, gene hits from each cell type were filtered to include only genes with adjusted p-value < 0.05 and absolute LogFC > 0.25. Enrichment testing was conducted separately for upregulated and downregulated gene sets using the *enrichGO* function, with background genes defined as all genes tested for differential expression in the corresponding analysis. Multiple testing correction was applied using the Benjamini-Hochberg method, and GO terms with adjusted p-values < 0.05 were considered significant.

#### NeuronChat

Inferred interactions were identified via the mouse database of receptor-ligand pairs and the SCT data layer of the snRNAseq data. Cord diagrams show both directionality of interaction as indicated by arrow direction, and the relative strength of that interaction to the others in the network via arrow line weight. Aggregation across pairs was performed using the “weight” parameter. Cell-cell communication strength is calculated as the product of the ligand abundance of a “sender” cell group and the target abundance of a “receiver” cell group. Significance is determined by permutation test where group labels of cells are randomly permuted and the communications strength is recalculated. Connection counts refer to the number of non-zero sender-receiver interactions amongst cell groups. For sample level differential analysis, NeuronChat^97^ was run individually on each sample and the resulting metrics were aggregate using mean where applicable or the replicate level data is shown. Differential strength was calculated via subtractive analysis of group means, with positive values indicating increased strength in the *Chd8^+/-^* group. Signaling supertypes (e.g. Glut) represent the mean aggregation of individual interaction pairs using that ligand. Information flow refers to a weighted product incorporating number of interactions and their strengths. Output figures generated summarizing cell-cell communication were generated using NeuronChat and R.

#### Calcium Imaging - Stereotactic injections

We utilized adult mice of either sex housed and bred in the UCSF animal facility. To image prefrontal activity in acute slices from wildtype or mutant animals, mice were injected with AAV5.hSyn.GCaMP6f.WPRE.SV40 (Penn Viral Core). Mice were anesthetized with 2% isoflurane and mounted in a stereotactic frame. Craniotomies were made according to stereotaxic coordinates relative to Bregma. Coordinates for injection into mPFC were (in mm relative to Bregma): +1.7 anterior– posterior (AP), –0.3 mediolateral (ML) and –2.75 dorsoventral (DV). Imaging was performed 4-6 weeks later.

#### Slice preparation and imaging

300 µm coronal slices were prepared as previously described^98^. Immediately after preparation slices were transferred to an N-Methyl-D-Glucamine (NMDG)-based recovery solution for 10 minutes before being transferred to ACSF for the remainder of their recovery. The NMDG-based solution was maintained at 32° C, and consisted of the following (in mM): 93 N-Methyl-D-Glucamine (NMDG), 93 HCl, 2.5 KCl, 1.2 NaH2PO4, 30 NaHCO3, 25 glucose, 20 HEPES, 5 Na-ascorbate, 5 Na-pyruvate, 2 thiourea, 10 magnesium sulfate, 0.5 calcium chloride. ACSF contained the following (in mM): 126 NaCl, 26 NaHCO3, 2.5 KCl, 1.25 NaH2PO4, 1 MgCl2, 2 CaCl, and 10 glucose. For imaging slices were perfused with ‘Active’ ACSF which was identical to normal ACSF except containing elevated KCl (3.5 mM vs 2 mM) and reduced CaCl (1.2 mM vs 2 mM). Slice data was acquired at 10 Hz on an Olympus BX51 upright microscope with a 20× 1.0NA water immersion lens, 0.5× reducer (Olympus), and ORCA-ER CCD Camera (Hamamatsu Photonics). Illumination was delivered using a Lambda DG4 arc lamp (Sutter Instruments). Light was delivered through a 472/30 excitation filter, 495nm single band dichroic, and 496nm long pass emission filter (Semrock). All movies consisted of 36000 frames acquired at 10Hz (1 hr) with 4×4 sensor binning yielding a final resolution of 256 × 312 pixels. Light power during imaging was 100 – 500 μW/mm2. The Micro Manager software suite (v1.4, NIH) was used to control all camera parameters and acquire movies. When necessary, drift correction was conducted in MATLAB using custom written code^99^ modified to utilize individual neurons as fiducials, whose positions were averaged and aligned every 5 seconds. We segmented neuronal signals using a modified PCA/ICA approach^100^ modified as previously described^101^ so that each segment was expressed as a binary ROI consisting of pixels representing a single neuron. To deconvolve neuronal signals from background neuropil signals, we subtracted the mean signal from each identified segment from the mean value in pixels surrounding the edge of the segment (we excluded pixels that belonged to another ROI). Signals were subsequently lowpass filtered to remove high frequency noise using the Matlab command: designfilt(’lowpassfir’, ‘PassbandFrequency’, 0.5, ‘StopbandFrequency’, .65, ‘PassbandRipple’, 1, ‘StopbandAttenuation’, 25). We then detected events corresponding to epochs in which each neuron was active using threshold-based event detection as previously described4. We detected increases in (F-F0)/F0 exceeding 2.5σ over one frame, and then further thresholded these events by keeping only those events which exceeded a 8σ increase over baseline and had an integrated area of 200 σ. σ is the standard deviation of (F-F0)/F0. All further analysis was performed on binary rasters of detected events for each slice.

#### Cranial Window Implantation Surgery

Cranial window implantation in adult mice (6–8 weeks old) was performed as previously described^102^. Briefly, the mouse was anaesthetized by gaseous isoflurane (4% for induction, 1.5-2% for maintenance) delivered through a vaporizer system and mounted on a stereotaxic frame. Throughout the surgery the mouse was kept warm with a heating pad. Ophthalmic ointment was applied to prevent eye desiccation and irritation. Carprofen (5 mg/kg, i.p.), buprenorphine (0.1 mg/kg, s.c.), enrofloxacin (5 mg/kg, s.c.), and dexamethasone (2 mg/kg, intramuscular) were administered. The fur over the surgical site was shaved, and the scalp was cleaned with 3 rounds of alternating 70% ethanol and iodine solution (Betadine®). A midline scalp incision was made, and the periosteum was gently scraped off from the skull using a scalpel. A circular piece of the skull (centered at AP = –1 mm, ML = 1.5 mm) was removed with a trephine (diameter = 2.3 mm, Fine Science Tools, Foster City, CA, USA) driven by a high-speed micro-drill (Foredom K1070, Blackstone Industries, LLC, Bethal, CT, USA). The imaging port was made by gluing a circular cover glass (#2, diameter = 2.2 mm) underneath a donut-shaped glass (#1, inner diameter = 2 mm, outer diameter = 3 mm; Potomac Photonics, Inc., Baltimore, MD, USA). The imaging port was mounted so that the bottom cover glass fit snugly into the cranial window and the top glass donut rested above the skull. The imaging port was secured with a UV-cured adhesive (Fusion Flo, Prevest DenPro, Jammu, India) onto the skull. After the solidification of the adhesive, the scalp flaps were closed with suture. After 2 weeks of recovery, the central piece of the scalp was excised, the edge of the scalp flap was secured to the tissue underneath with cyanoacrylate (Vetbond), and a custom-made stainless-steel head-bar was secured over the skull with dental cement (Jet Denture Repair, Lang Dental, Wheeling, IL, USA). The mouse received enrofloxacin, buprenorphine, and carprofen once per day for two extra days post-surgery and was allowed to recover for an additional week prior to imaging.

#### Thin skull preparation

The thin skull procedure was performed on adolescent (1 month old) mice as previously described^103^. Briefly, the mouse was anesthetized with a mixture of ketamine (20 mg/ml) and xylazine (2.0 mg/ml) in 0.9% sterile saline administered intraperitoneally (5 ml/kg body weight). Ophthalmic ointment was applied to the eyes to prevent desiccation and irritation, and the fur over the scalp was removed with a blade. A midline incision was made through the scalp and the periosteum was gently scraped off from the skull. A high-speed micro-drill (Foredom K1070, Blackstone Industries, LLC, Bethal, CT, USA) and a microblade were used to thin a small region of the skull to ∼20 μm thickness. A custom-made head-plate with a central opening was attached to the skull by cyanoacrylate glue (Krazy Glue, Elmer’s Products, Westerville, OH, USA), centered over the thinned region. The head-plate was secured onto a custom-made metal baseplate to stabilize the mouse’s head during imaging. Two-photon imaging was performed as described above. After imaging, the head-plate was detached from the skull, the skull was cleaned with sterile saline, and the scalp was sutured. The same region was imaged again 7 days later.

#### In vivo two-photon (2P) imaging and analysis

2P imaging of dendritic spines was performed on a 2P microscope (Ultima IV, Bruker Co., Middleton, WI) equipped with a 40× NA = 0.8 water immersion objective (LUMPlanFl/IR, Olympus, Japan) and an ultrafast 2P laser (Mai Tai, Spectra-Physics, Santa Clara, CA) operating at 940 nm. The mouse was anaesthetized with an intraperitoneal injection of a mixture of 85 mg/kg ketamine and 8.5 mg/kg xylazine in 0.9% saline and mounted on a custom-made stage for imaging. Stacks of images were acquired with a Z-step size of 1 µm at 4X zoom. Relocation of the same dendrites in subsequent imaging sessions was achieved by reference to blood vessels and the dendritic branching pattern. The same dendrites were imaged again 1 week after the first imaging session.

Data analysis was performed in ImageJ as previously described^103^. Typically, 150–200 spines were analyzed per animal per session. The percentage of spines formed/eliminated was calculated as the number of spines formed/eliminated divided by the total number of spines counted from the previous imaging session. Dendritic spine density was calculated as the number of dendritic spines per 10 µm of dendritic segment length on the first imaging day.

#### EEG/EMG recording and analysis

The EEG/EMG implant consists of a mini connector (Samtec Inc., SFMC-102-01-S-D) soldered to 2 EEG and 2 EMG electrodes made from Teflon-coated, multi-filament stainless steel wires (0.002” in diameter, Medwire 316 SS7/44T, Sigmund Cohn Corp.). The mouse was anaesthetized by gaseous isoflurane (4% for induction, 1.5-2% for maintenance) delivered through a vaporizer system and mounted on a stereotaxic frame. Throughout the surgery the mouse was kept warm with a heating pad. Ophthalmic ointment was applied to prevent eye desiccation and irritation. Buprenorphine (0.1 mg/kg, s.c.), carprofen (5 mg/kg, i.p.), and enrofloxacin (5 mg/kg, s.c.) were injected. The fur over the surgical site was shaved, and the scalp was cleaned with 3 rounds of alternating 70% ethanol and iodine solution (Betadine®). A piece of the scalp was excised with scissors to expose the skull. The edge of the scalp flap was secured to the tissue underneath with cyanoacrylate (Vetbond). The exposed skull surface was gently scraped with a scalpel to remove the periosteum. Small holes (one at AP = + 2 mm, ML = 2 mm; the other at AP = -3 mm, ML = 2 mm) were drilled through the skull using a high speed microdrill (Foredom K.1070) with fine drill bits. The EEG electrodes were inserted into the holes, positioned between the skull and the brain surface, and secured with a UV-curable adhesive (Fusion Flo, Prevest DenPro). The EMG electrodes were positioned under the nuchal trapezoid muscles and secured with Vetbond. The exposed skull surface was covered with a thin layer of Vetbond. The EEG/EMG implant connector was then further secured onto the skull by dental cement (Jet Denture Repair, Lang Dental). Enrofloxacin (5 mg/kg, s.c.) and carprofen (5 mg/kg, i.p.) were given at 24 h after surgery. The mouse was allowed to recover for 7 days before EEG/EMG recording starts. A BL-420N biological signal acquisition and processing system (Chengdu Techman, China) was used for EEG/EMG recording. EEG was sampled at 200 Hz and band-pass filtered between 1-100 Hz; EMG was sampled at 200 Hz and band-pass filtered between 1-2K Hz. Recorded data was annotated manually and subsequently processed using custom-written Matlab programs (R2022, MathWorks Inc.).

---

**Figure S1:**
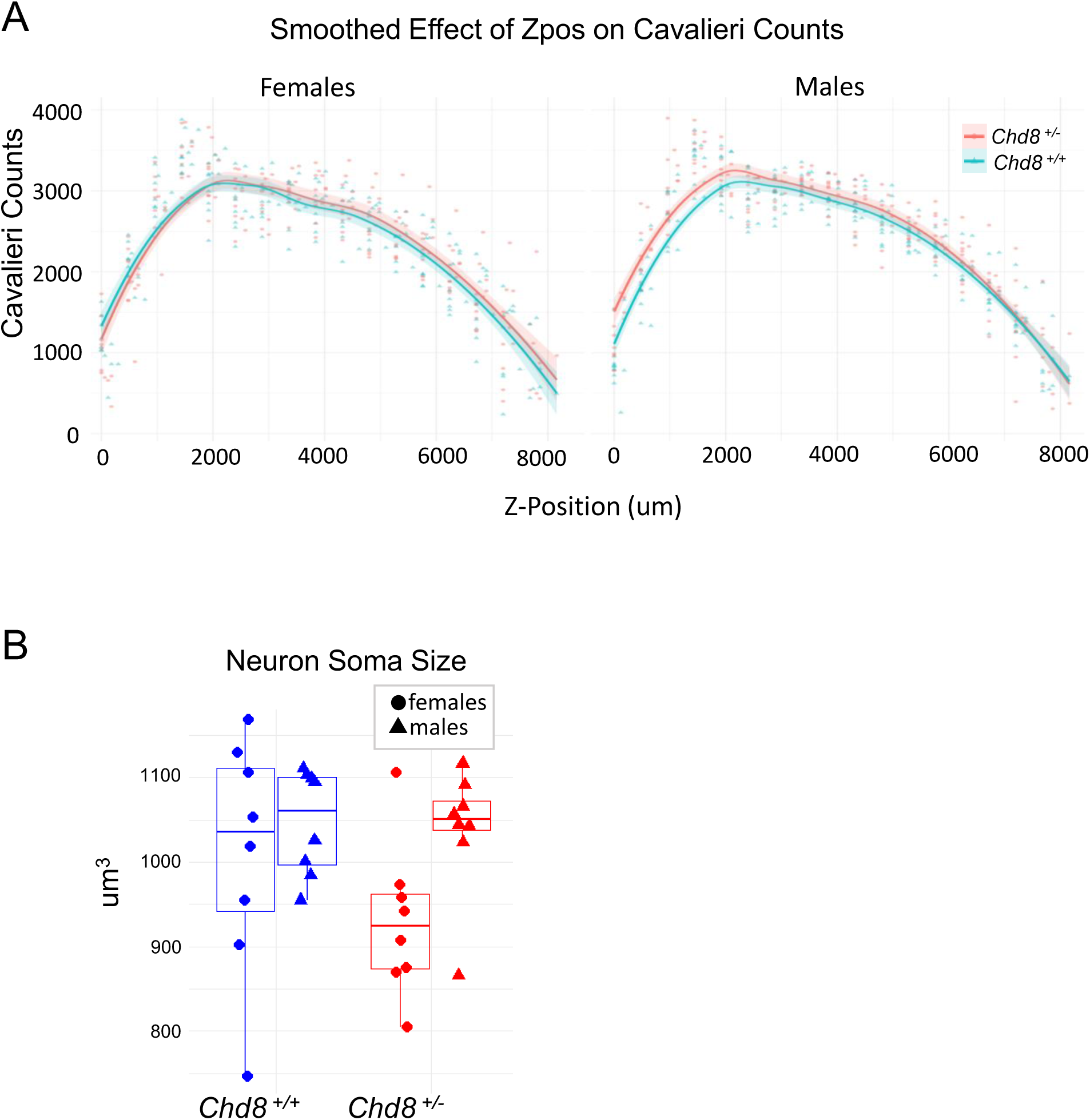
A) Cortical volume was quantified across rostral–caudal section in *Chd8^+/−^* and *Chd8^+/+^* mice using the Cavalieri method. In male *Chd8^+/−^* mice, increased volume was observed predominantly in more rostral regions, with clearer separation of genotype averages along the anterior Z-positions. Shaded portions of the plots indicate the more caudal regions (0–3000 µm), where volume differences are most evident. B) Neuronal soma size, measured by unbiased stereology in the same rostro-caudal range, did not differ significantly between genotypes. Data are stratified by sex and genotype. Error bars represent SEM. Statistical comparisons were performed using two-way ANOVA (unstratified: P = 0.23; males: P = 0.81; females: P = 0.19).

**Figure S2:**
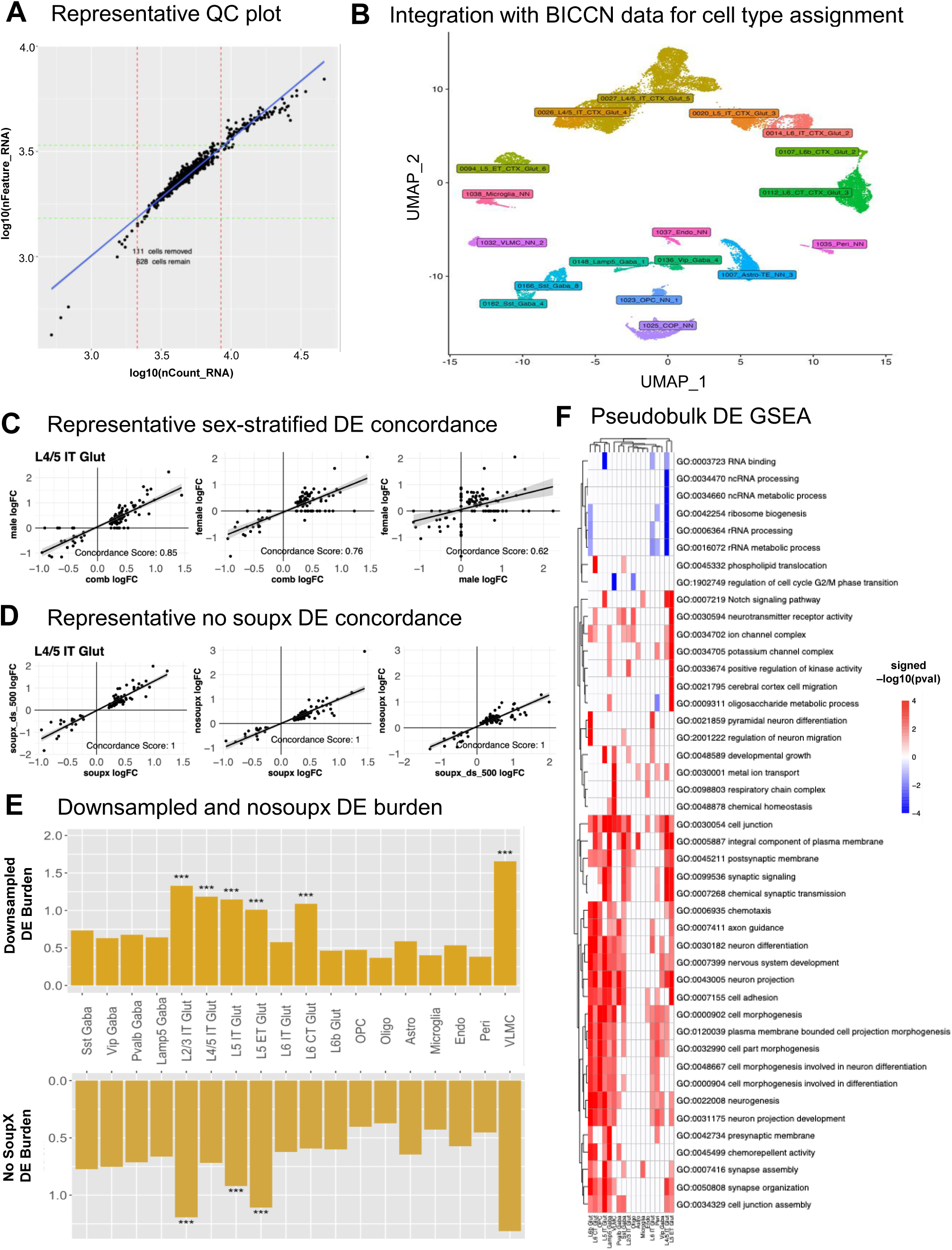
A) Scatterplot of log10-transformed total RNA counts (nCount_RNA) versus detected genes (nFeature_RNA) per cell in one batch. Custom MAD-based thresholds were chosen per batch to filter low-quality cells. Red and green dotted lines indicate nCount_RNA and nFeature_RNA cutoffs, respectively; cells outside these thresholds were excluded from downstream analysis. B) UMAP showing all *Chd8* mutant and wild-type nuclei integrated with BICCN data and annotated with predicted BICCN cell type labels, which are consistent with cell type identities derived from our independent annotation pipeline. C) Scatterplots comparing log fold changes (logFC) of the top 100 DEGs identified with full SoupX, non-downsampled DE analysis (“soupx”) to their logFCs in the no-SoupX (“nosoupx”) and downsampled to 500 nuclei + SoupX (“soupx_ds_500”) analyses for a representative cell type. Concordance scores were calculated as the proportion of genes with the same direction of change between methods. D) Scatterplots comparing logFC of the top 100 DEGs identified with male-and-female-combined (“comb”) DE analysis to their logFCs in the male and female DE analyses for a representative cell type. E) Bar plots of DEG burden across cell types (Burden = DE genes / total genes expressed * 100) for downsampled (top) and no-SoupX (bottom) DE analyses. A permutation test with 10,000 iterations was used to assess statistical significance and confirmed that the observed trends were similar to those in the full SoupX, non-downsampled analysis. F) Heatmap of normalized enrichment scores (NES) for a curated set of biologically relevant GO terms identified by GSEA performed on pseudobulk DEGs for each cell type.

**Figure S3:**
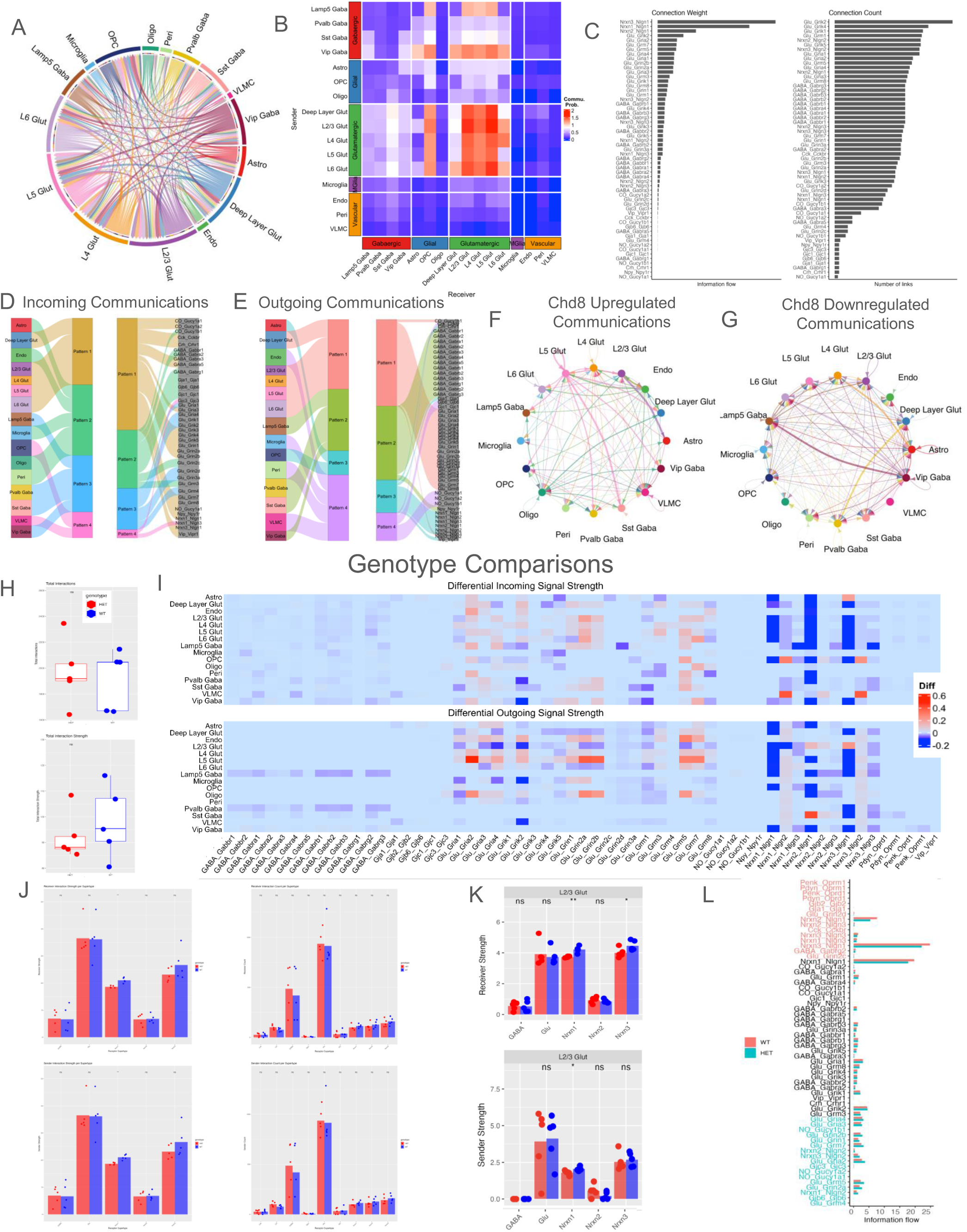
A) Overview of all inferred connections by cell type shows generally balanced co-directional signaling for each cell type. B) The communication probability is strongest within Glutamatergic cell types and between Glutamatergic and GABAergic populations. C) The strength and count of connections anchored by the listed receptor highlight networking drivers. D, E) Cell types are grouped based on the similarity of their incoming or outgoing connections showing similarities correlated with similar cell identities and neurotransmitter types. F, G) Circle plots showing the upregulated (F) and downregulated (G) intercellular communications in *Chd8^+/-^* compared to control. The width of each link indicates the absolute difference between Chd8 and control, in the sum of communication strength values over all interaction pairs. H) The relative strength (top) of all network connections is similar in both WT and *Chd8^+/-^* (“HET”) animals but the number of connections (bottom) shows some elevation in the *Chd8^+/-^* animals. I) Heatmap showing the differential outgoing (upper panel) and incoming (lower panel) signal strength between *Chd8^+/-^* and control. Given the cell group and interaction pair, the outgoing (or incoming) signal strength is defined as the sum of communication strength over the links from (or to) the cell group. The color bar indicates the difference in the outgoing (or incoming) signal strength values between *Chd8^+/-^* and control. Interaction pairs with signal strength of 0 for all cell types were omitted from plotting. J) Replicate level quantification of receptor supertype sender (top) and receiver (bottom) differences shows dysregulation only in the Nrxn1 strength metrics on the level of the entire dataset. K) Further cell type specific inspection indicates that the L2/3 population is the only significantly affected population driving the dysregulation. L) Bar chart comparing the information flow between *Chd8^+/-^* and control for each interaction pair at the full dataset level. While the HET samples show both decreased and increased information flow in some interaction pairs, those that are increased have a vastly greater magnitude thus dominating the effects. In contrast, the increased information flow seen in the glutamate interactions is present in connections with decreased overall magnitude but is considerably consistent across each glutamate-receptor-ligand pair and reveals the glutamate network dominating the rankings of information flow increases in the HET.

**Figure S4:**
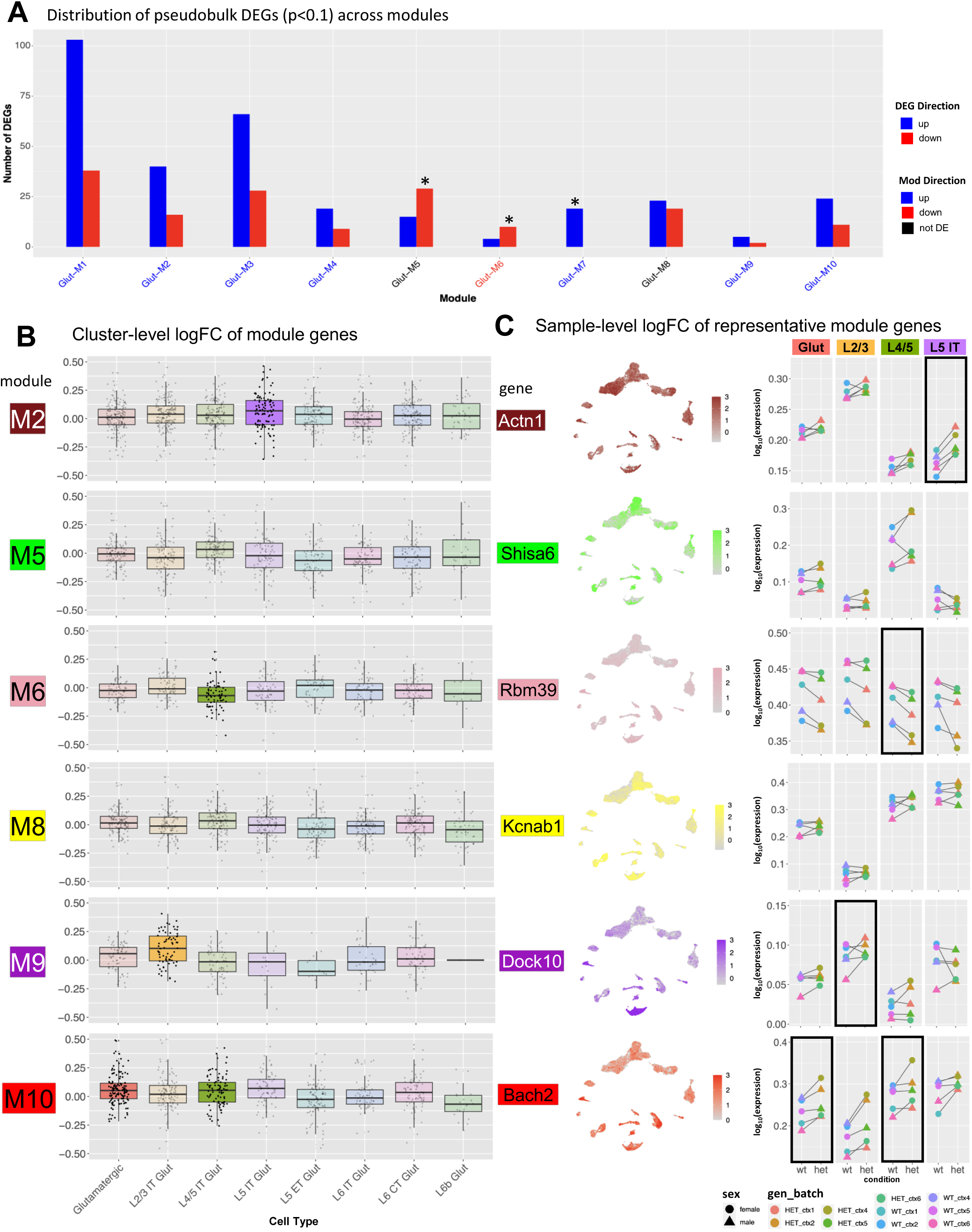
A) Bar plot showing the distribution of DEGs (p < 0.1) across glutamatergic modules. Bars indicate the number of genes upregulated (blue) or downregulated (red) in at least one cell type per module. Modules Glut-M5 and Glut-M6 showed significant enrichment for downregulated DEGs (Fisher’s exact test, FDR-adjusted *p < 0.05), while Glut-M7 showed significant enrichment for upregulated DEGs. Overall, the direction of module-level change tends to agree with the predominant direction of DE for the genes within each module. B) Box plots for the remaining glutamatergic modules, showing logFCs of module genes across cell types. Opaque boxes indicate subtypes where the module is significantly differentially expressed (Student’s t-test comparing mean logFC of each module to grey module, Bonferroni-adjusted p < 0.05). C) Feature plots (left) show relative expression of representative genes from each module in UMAP space. Dot plots (right) display median sample-level expression of the same genes. Black boxes highlight subtypes where the module is differentially expressed or where consistent directional changes are observed across batches.

**Figure S5:**
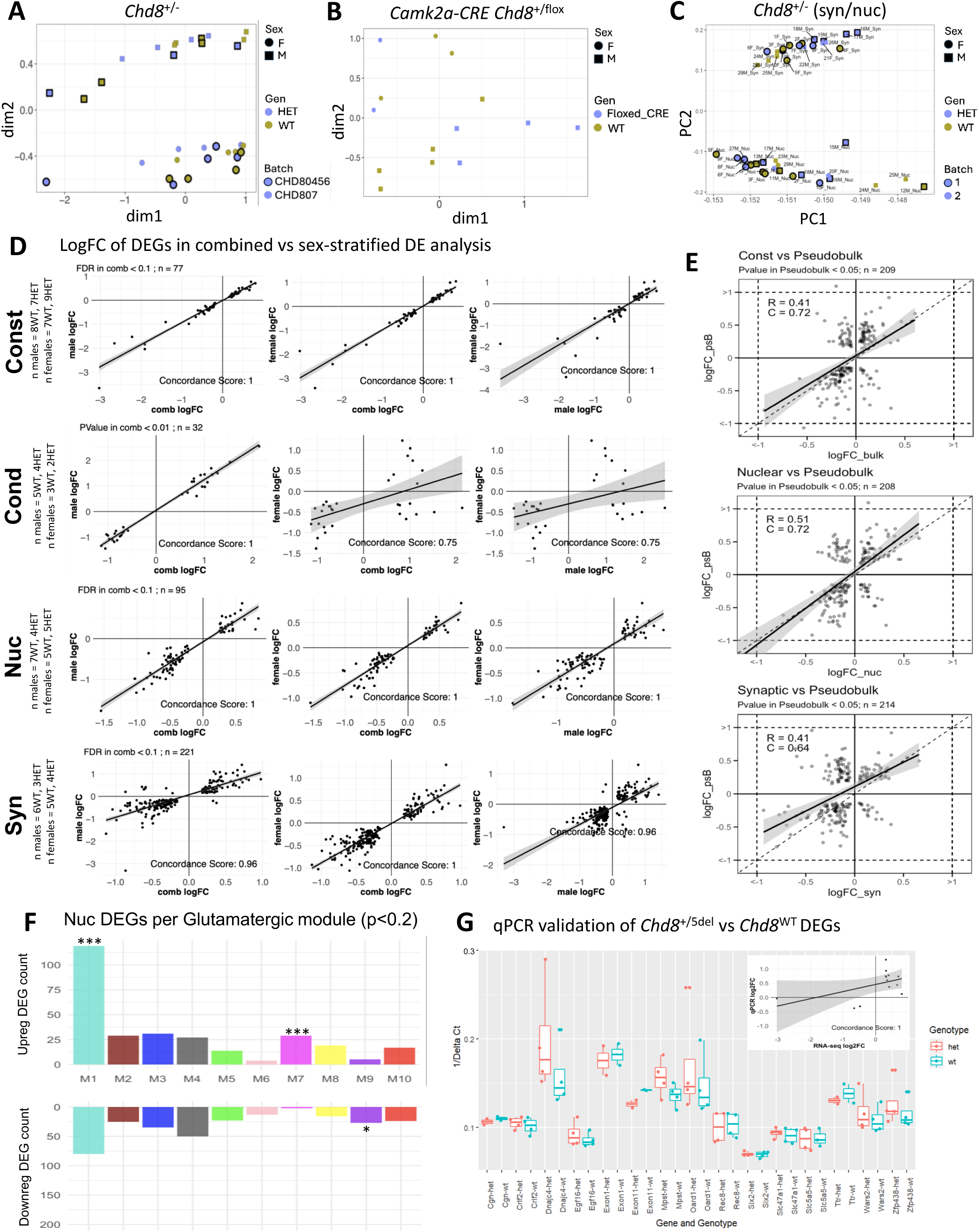
A-B) MDS plot of cortical *Chd8* mutant (A: *Chd8^+/-^*, B: *Camk2a-CRE Chd8^+/flox^*) and WT samples, with the the second dimension (“dim2”) reflecting biological sex. C) PCA plot of cortical *Chd8^+/-^* and *Chd8^WT^* synaptic and nuclear samples, where leading dimensions distinguish sample fraction. D) Log fold change (logFC) concordance scatterplots comparing logFC of DEGs from the combined (”comb”) analysis to their logFCs in each of the sex-stratified analyses (“male”, “female”). Only genes passing FDR < 0.1 or P < 0.01 in combined datasets were used for comparisons. Each row represents one of 4 bulk RNA-seq datasets: constitutive *Chd8^+/-^* (“Const”), conditional *Camk2a-CRE Chd8^+/flox^* (“Cond”), nuclear fraction of *Chd8^+/-^* (“Nuc”), and synaptic fraction of *Chd8^+/-^* (“Syn”). Plots display gene-level logFC correlations between pairs of analyses with linear regression fits and concordance scores indicating the fraction of genes regulated in the same direction. E) LogFC scatterplots comparing constitutive *Chd8^+/-^*(“Const”) to pseudobulk snRNA-seq DEGs (top), nuclear fraction *Chd8^+/-^*(“Nuc”) to pseudobulk DEGs (middle), and synaptic fraction *Chd8^+/-^*(“Syn”) to pseudobulk DEGs. Genes were selected based on significance in pseudobulk analysis (P < 0.05) and differential expression in each respective bulk RNA-seq analysis. Pearson correlation coefficients (R) and concordance scores (C) are shown on each plot. F) Bar plots showing counts of nuclear differentially expressed genes (DEGs) per module, separated into upregulated (top) and downregulated (bottom). Enrichment was assessed using Fisher’s exact test, comparing the proportion of up- vs. downregulated DEGs in each module relative to all other modules, with multiple testing correction (Benjamini-Hochberg). M1 and M7 were significantly enriched for upregulated DEGs (p*** < 0.001), while M9 was enriched for downregulated DEGs (p* < 0.05). G) Box plots showing the median 1/ΔCt (relative to GAPDH) across biological replicates for 13 constitutive *Chd8^+/-^* bulk RNA-seq DEGs. *Chd8* Exons 1 and 11 (“Exon1”, “Exon 11”) were included as positive controls. A Wilcoxon text showed nonsignificant differences in 1/ΔCt between *Chd8* mutants and wild-types, all 13 genes tested showed the same direction of change as RNA-seq DE analysis. Inset shows scatterplot of RNA-seq logFC versus qPCR logFC, with concordance calculated as in (D-E).

